# Mitochondrial genome sequencing and analysis of the invasive *Microstegium vimineum*: a resource for systematics, invasion history, and management

**DOI:** 10.1101/2023.02.10.527995

**Authors:** Craig F. Barrett, Dhanushya Ramachandran, Chih-Hui Chen, Cameron W. Corbett, Cynthia D. Huebner, Brandon T. Sinn, Wen-Bin Yu, Kenji Suetsugu

**Author notes:** Corresponding author information: Craig F. Barrett, ORCiD 0000-0001-8870-3672.

## Abstract

**Premise of the Research:** Plants remain underrepresented among species with sequenced mitochondrial genomes (mitogenomes), due to the difficulty in assembly with short-read technology. Invasive species lag behind crops and other economically important species in this respect, representing a lack of tools for management and land conservation efforts.

**Methodology:** The mitogenome of *Microstegium vimineum*, one of the most damaging invasive plant species in North America, was sequenced and analyzed using long-read data, providing a resource for biologists and managers. We conducted analyses of genome content, phylogenomic analyses among grasses and relatives based on mitochondrial coding regions, and an analysis of mitochondrial single nucleotide polymorphism in this invasive grass species.

**Pivotal Results:** The assembly is 478,010 bp in length and characterized by two large, inverted repeats, and a large, direct repeat. However, the genome could not be circularized, arguing against a “master circle” structure. Long-read assemblies with data subsets revealed several alternative genomic conformations, predominantly associated with large repeats. Plastid-like sequences comprise 2.4% of the genome, with further evidence of Class I and Class II transposable element-like sequences. Phylogenetic analysis placed *M. vimineum* with other *Microstegium* species, excluding *M. nudum*, but with weak support. Analysis of polymorphic sites across 112 accessions of *M. vimineum* from the native and invasive ranges revealed a complex invasion history.

**Conclusions:** We present an in-depth analysis of mitogenome structure, content, phylogenetic relationships, and range-wide genomic variation in *M. vimineum’s* invasive US range. The mitogenome of *M. vimineum* is typical of other andropogonoid grasses, yet mitochondrial sequence variation across the invasive and native ranges is extensive. Our findings suggest multiple introductions to the US over the last century, with subsequent spread, secondary contact, long-distance dispersal, and possibly post-invasion selection on awn phenotypes. Efforts to produce genomic resources for invasive species, including sequenced mitochondrial genomes, will continue to provide tools for their effective management, and to help predict and prevent future invasions.

## Introduction

Invasive species cause damage to natural, agricultural, and urban ecosystems, equating to billions of dollars (USD) in economic and environmental loss (Pimentel et al., 2005; Simberloff et al., 2013). Such problems have been exacerbated by climate change and greater interconnectedness across the globe (Finch et al., 2021). Genomic resources provide practitioners and researchers with a baseline of powerful tools in medicine, agriculture, and virtually all areas of the life sciences, yet such tools are generally lacking for invasive species compared to those in crop and animal systems (Matheson and McGaughran, 2022). However, the widespread availability and increasing affordability of genome sequencing technologies and bioinformatic platforms are changing the landscape of invasion biology (North et al., 2021). For example, such advances in genomics are allowing more nuanced reconstructions of invasion history (van Boheemen et al., 2017; Sutherland et al., 2021; Bieker et al., 2022), linking of genotypic and phenotypic variation (Turner et al., 2021; Revolinski et al., 2022), epigenetics (Banerjee et al., 2019; Mounger et al., 2021), and forecasting of potential future invasions (Hudson et al., 2021).

Generally speaking, plant mitochondrial genomes (‘mitogenomes’) have experienced less attention than plastid or nuclear genomes (Mower et al., 2012). This is largely due to their extensive variability in structural dynamics and repetitive DNA content, making them difficult targets for complete genomic sequencing (Palmer and Herbon, 1988; Alverson et al., 2010). This is in contrast to animal mitogenomes, which evolve rapidly in terms of substitution rates but are more structurally conserved. In combination with their smaller size (10-20 kb in animals vs. 100 kb to > 10 Mb in plants; Gualberto et al., 2014), animal mitogenome sequencing is more straightforward than in plants, making animal mitogenomes significantly better represented across the Tree of Life. Improvements in long read sequencing technology, however, have rekindled interest in plant mitochondrial genomics, allowing the assembly of complete or nearly complete mitogenomes, which often display repetitive regions and structural isoforms making them difficult or impossible to assemble with short-read technologies (Kovar et al., 2018; Jackman et al., 2020). Analyses of plant mitogenomes have revealed an array of structures, including circular genomes, “master circles” with sub-stoichiometric circular structures, linear structures, multi-chromosomal structures, and branched structures (Bendich, 1993; Sloan, 2013; Wu et al., 2015; 2022).

Plant mitogenomes typically contain 50-60 genes, including those encoding protein products (CDS, or coding DNA sequences), transfer RNAs (tRNAs), and ribosomal RNAs (rRNAs) (Gualberto et al., 2014). They are also known to contain plastid-like regions, likely as remnants of both ancient and recent intergenomic transfers and gene conversion events, representing up to 10.3% of the mitogenome in the date palm *Phoenix dactylifera* (Fang et al., 2012). Additionally, plant mitogenomes have been demonstrated to house foreign DNA, possibly remnants of ancient or more recent close biotic interactions (e.g. Rice et al., 2013; Sanchez-Puerta et al., 2019; Sinn and Barrett, 2020; Lin et al., 2022).

Grasses are overrepresented in terms of complete mitogenomes among plant families, with 67 complete genomes in NCBI GenBank, though more than half of these comprise multiple accessions of a few crop species (e.g. *Hordeum vulgare, Oryza sativa, Triticum aestivum, Zea mays*; last accessed 18 November, 2022). However, grasses are also overrepresented among invasive plant species (Daehler, 1998; Kerns et al., 2020), allowing for meaningful comparisons among invasive and non-invasive species within this ecologically and economically important family. Only a handful of mitogenomes have been sequenced for invasive plants, and most are grasses [e.g. *Silene vulgaris* (Caryophyllaceae); *Chrysopogon zizianoides*, *Coix lacryma-jobi*, *Eleusine indica*, and *Lolium perenne* (Poaceae)]. Thus, studies of mitochondrial genome dynamics in invasive plants, and potential applications in their effective control, are in their infancy. For example, a simulation study by Hodgins et al. (2009) explored the possibility of incorporating cytoplasmic (mitochondrial) male sterility alleles in the control of invasive plants by limiting pollen production. Yet, empirical data and sequenced reference mitogenomes are too few to test the effectiveness of such approaches more broadly in invasive plant species, nearly all of which can be categorized as non-model species.

*Microstegium vimineum* (stiltgrass) is an aggressive, established invader of eastern North American forest ecosystems, and is one of the most damaging invasive species on the continent (e.g. Huebner, 2010a; Johnson et al., 2015). Likely introduced as packing material for porcelain in the early 1900s (Fairbrothers and Gray, 1972), this species has spread to 30 US states, and is expanding into Canada, the northeastern US, and the northern US Midwest (Huebner, 2010a; 2010b; Mortensen et al., 2009; Rauschert et al., 2010; Barrett et al., 2022). Further, it is hypothesized that *M. vimineum* was introduced multiple times in the US, first in the southeastern US, and later in the Northeast, with subsequent spread and secondary contact, providing an apt case study in the genomic dynamics of the invasion process (Novy et al., 2013; Barrett et al., 2022). Recently published plastid and nuclear genomes are now available for this species (Welker et al., 2020; Ramachandran et al., 2021, respectively), but a complete mitogenome is lacking. Therefore, the objective of this study is to assemble a reference mitogenome for *M. vimineum*, with the goal of aiding studies of invasion history, evolution, ecology, and management. We explore genome structure and content, phylogenetic relationships of *Microstegium*, and patterns of mitogenomic variation across the native and invasive ranges with respect to invasion history in *M. vimineum*.

## Materials and Methods

### Organellar genome sequencing and assembly

Leaf material was sampled from a growth chamber-grown accession (seed from Potomac Ranger District, Monongahela National Forest, West Virginia, USA), flash-frozen in liquid Nitrogen, and stored at −80C. DNAs/RNAs were extracted, and PacBio (DNA) and Illumina (DNA and RNA) sequencing were conducted as described in Ramachandran et al. (2021). The software seqtk v.1.0-r31 (https://github.com/lh3/seqtk) was used to randomly subsample PacBio reads (400,000 reads). MegaBLAST from the NCBI BLAST+ suite (Camacho et al., 2008) was conducted with the subsampled read pools against the mitochondrial genome of *Sorghum bicolor* (NCBI GenBank number NC_008360) in Geneious v.10.0.9 (http://www.geneious.com/), specifying an e-value of 1e^-5^, and binning into ‘hits’ vs. ‘no hits,’ keeping only reads >20 kb in length. The resulting positive BLAST hits for each set were then assembled with CANU v.2.2 under default parameters (Koren et al., 2017). The resulting graphs from CANU (.gfa files) were inspected in BANDAGE v.0.9.0 (Wick et al., 2015) to visualize contiguity and coverage of the assemblies. Resulting scaffolds were further assembled into a single scaffold in Geneious using the native overlap-layout-consensus ‘*de novo* assembly’ option. Mitochondrial and plastid contigs were identified using the live annotation feature in Geneious, with the annotations from *Sorghum bicolor* (mitochondrial), and an accession of *M. vimineum* (plastome; accession TK124, GenBank number MT610045) at a 70% threshold, respectively. Circlator was used to attempt to circularize the scaffold (Hunt et al., 2015).

Mitochondrial and plastid contigs were extracted separately as FASTA files. FLYE was then used to correct the mitochondrial and plastid scaffolds with ten polishing iterations using the PacBio data (Kolmogorov et al., 2019). The assembly was further polished with Illumina data using PILON (Walker et al., 2014). Illumina data (8,605,412 read pairs from accession WV-PRD-2-4, the same collection used for PacBio sequencing) were trimmed with BBDUK v.38.51 (https://sourceforge.net/projects/bbmap) to remove Illumina adapters, low quality bases (minimum quality = 6), and low-complexity regions (minimum entropy = 0.5, maximum GC content = 0.9). Illumina reads were then mapped to the organellar assemblies with NGM (Sedlazeck et al., 2013) to output a .sam alignment file. The .sam file was then sorted and indexed with SAMTOOLS v.1.7 (Li et al., 2009). The PacBio assemblies and sorted .bam file were then used for error correction/polishing with PILON.

The resulting polished FASTA file was imported into Geneious and annotated using the ‘live annotation’ feature, at a 75% similarity threshold, using the mitochondrial annotations from *Coix lacryma-jobi* var. *ma yuen* (GenBank accession number MT471100), *Sorghum bicolor* (same accession as above), *Oryza sativa* (ON854123), *Zea mays* (CM025451), and *Saccharum officinarum* (MG969496) and the plastid annotation from *Microstegium vimineum* (same as above). Annotations were then checked visually to confirm proper start/stop codons and to investigate the presence of premature stop codons, suggesting mis-annotations. The annotation was exported from Geneious as a GenBank Flat File and converted to a feature table with GB2Sequin (Lehwark and Greiner, 2019) via the ChloroBox portal (https://chlorobox.mpimp-golm.mpg.de). The resulting feature table was downloaded and manually edited to ensure the correct orientation of exons in genes containing them. The annotation (feature table + fasta file) was then submitted to GenBank though the BankIt web portal (https://www.ncbi.nlm.nih.gov/WebSub).

### Analyses of repetitive DNA, plastid-like DNA, RNA editing, and structural variation

Geneious was used to identify large, identical repeats >1,000 bp, using the native Repeat Finder plugin (https://www.geneious.com/plugins/repeat-finder/) as well as the self-dotplot function, with a window size of 500 bp and tile size of 100 kb. REPuter (Kurtz et al., 2001; via https://bibiserv.cebitec.uni-bielefeld.de/reputer/) was further used to detect repetitive regions >8 bp in length in forward, reverse, reverse-complement, and palindromic configurations (edit and Hamming distances = 0). Plastid-like regions were identified by annotating the mitogenome with all plastid genes from *M. vimineum* (Genbank # MT610045) in Geneious at a 60% similarity threshold, in order to detect degraded or pseudogenized plastid-like sequences. Further, to identify plastid-like regions not corresponding to annotated genes, Illumina reads from accession WV-PRD-2-4 (same as above) were mapped to the reference mitogenome to identify putative plastid-like regions with higher than expected coverage depth. These regions were annotated in Geneious as having >3x standard deviations in coverage depth relative to the rest of the genome.

RNAseq reads from the same collection from young, developing leaf tissue (NCBI Sequence Read Archive accession SRX12501806), were mapped to the reference genome using the Geneious read mapper for RNAseq data, and SNPs were called to identify putative RNA editing sites for all CDS (e.g. C → U). Minimum required coverage depth for a SNP was 10×, further requiring a minimum variant frequency of 0.9, such that only variants that differed among the RNAseq and DNAseq data were identified (i.e. the polished reference). Relative expression levels (transcripts per million, TPM) were calculated in two ways. First, all RNAseq reads were mapped to the plastome to filter plastid reads in Geneious, then the remaining reads were mapped to the mitochondrial annotation to quantify expression levels of all mitochondrial coding sequences (CDS). Then, the plastid-like region annotations were overlaid on the mitochondrial genomes, and plastid-filtered reads were mapped to assess whether plastid-like regions of the mitogenome displayed evidence of expression. All results were plotted in R with the packages dplyr v.1.0.10 (Wickham et al., 2023), ggplot2 v.3.3.6 (Wickham, 2016), and ggpubr v.0.4.0 (Kassambara, 2020). To investigate variation in mitogenome structure, eight random subsets of 200,000 PacBio reads were sampled with Seqtk and BLASTed against the *Sorghum* mitogenome (as above). BLAST hits were assembled with Flye, and the longest mitochondrial contigs were mapped to the reference genome model with the LASTZ v.1.04.22 (Harris, 2007) plugin for Geneious.

### Identification of transposable elements and foreign-aquired sequence

RepeatMasker version 4.1.1 (Smit et al., 2013) was used to discover and identify transposable elements (TEs) in the mitogenome assembly, usingRepBase-20181026 database of Viridiplantae (Bao et al., 2015) and a custom set of 1,279 *M. vimineum* consensus repeat sequences (Ramachandran et al., 2021). Additional repeat identification tools were used to screen for the presence of partial or truncated TE sequences in the mitogenome. HelitronScanner (Xiong et al., 2014) was used to identify helitrons using 5’and 3’ terminal motifs. Miniature inverted-repeat transposable elements (MITEs) were detected using the program MiteFinderII (Hu et al.,2018) under default settings.

Kraken 2 (v.2.1.2; Wood et al., 2019) was used to screen for the presence of interspecific genomic transfers, excluding those from the plastome. The mitogenome assembly, with repeats masked and plastid sequences removed, was decomposed into 100 bp segments using the reformat.sh script of the BBMAP suite (https://sourceforge.net/projects/bbmap/). The k-mers contained in the resulting 4,292 sequences were classified via screening against to precompiled Kraken 2 databases (available via: https://benlangmead.github.io/aws-indexes/k2): 1) PlusPFP database, which contained the complete genomes of plants, bacteria, archaea, viruses, fungi, human and UniVec vectors accessioned in NCBI’s RefSeq database; 2) sequences from 389 species in the Eukaryotic Pathogen, Vector and Host Informatics Resource Database (Amos et al., 2022). The NCBI Taxonomy (accessed 8 September 2022) was used for the annotation of classified sequences. The default values for k-mer length (35) and minimizer value (31) were used.

### Phylogenomic analyses using mitochondrial CDS

Mitochondrial genomes and mitochondrial protein coding sequences were downloaded from NCBI Genbank using the search terms ‘Poales,’ ‘mitochondrion,’ and ‘complete.’ *Cocos nucifera* (NC_031696) and *Phoenix dactylifera* (NC_016740) were chosen as outgroups, as both are members of the palm family (Arecales) and the ‘commelinid’ clade, to which Poales also belongs. Sequence annotations were extracted from Geneious and aligned with the codon-aware aligner MACSE v.2 (Ranwez et al., 2018). Alignments were then concatenated in Geneious, and sites with > 10% missing data were excluded (File S1). The final alignment was analyzed with maximum likelihood under the ‘GHOST’ heterotachy model (Crotty et al., 2020) in IQTree2 (Minh et al., 2020), which allows mixed substitution rates and branch lengths. This model is especially appropriate for lineages such as Poales, which have been shown in previous studies to exhibit heterotachy in phylogenomic estimates based on organellar DNA (e.g. Givnish et al., 2010; Barrett et al., 2016). This model approach avoids the need to partition the data by gene and codon, and can accommodate changes in substitution rates across branches and time. IQTree2 was run under the GHOST model with 1,000 ultrafast bootstrap pseudoreplicates (Hoang et al., 2018). The resulting tree file was visualized in FigTree v.1.4.4 (http://tree.bio.ed.ac.uk/) and edited with Adobe Illustrator v. 26.5 (Adobe, Inc., 2023).

### Assessment of relationships among mitochondrial haplotypes of M. vimineum

To characterize haploid single nucleotide polymorphisms (SNPs) in the mitogenome, we sampled 112 accessions from both the invasive (N = 74) and native ranges (Asia, n = 38), plus six accessions from different species of *Microstegium* as outgroups. Samples were either field collected in 2019-2020 (n = 66), or from herbarium specimens (n = 46) dating back to 1934 (Appendix A1). Total genomic DNAs were extracted via the CTAB method (Doyle and Doyle, 1987), and quantified via Qubit Broad Range DNA assay (Thermo Fisher Scientific, Waltham, Massachusetts, USA). DNAs were further visualized on a 1% agarose gel to assess degradation and diluted to 20 ng/ul with nanopure water. Illumina sequencing libraries were prepared with the SparQ DNA Frag and Library Kit at 2/5 volume (Quantabio, Beverly, Massachusetts, USA), which uses a fragmentase to shear genomic DNA, followed by end repair and adapter ligation. The shearing step for herbarium-derived DNAs was reduced to 1 min from 14 min, as these all showed some level of fragmentation prior to library preparation. Libraries were then amplified with primers matching the adapter sequences, adding dual-indexed barcodes (12 PCR cycles). Final library concentrations were determined via Qubit High Sensitivity DNA assay and pooled at equimolar ratios. Library pools were sequenced on two runs of 2 × 100 bp Illumina Nextseq2000 (v.3 chemistry) at the Marshall University Genomics core with samples from other studies, producing a total of ∼1 billion read pairs per run.

Reads were processed using a dedicated SNP calling pipeline (https://github.com/btsinn/ISSRseq; with scripts available at https://zenodo.org/record/5719146#.Y-EvfnbMKHs; Sinn, Simon et al., 2022). Briefly, reads were trimmed and filtered with BBDUK (http://sourceforge.net/projects/bbmap), with minimum read quality = PHRED 20, entropy and low complexity filters set to remove reads with < 0.1 or > 0.9 % GC content, kmer length set to 18, and the ‘mink’ flag set to 8. The reference genome was indexed and reads were mapped with BBMAP (https://sourceforge.net/projects/bbmap/). Here, plastid-like regions and one copy of each large repeat were removed from the reference genome to minimize drastic differences in coverage depth and plastid SNPs being misinterpreted as mitochondrial SNPs. The resulting .bam files were sorted and PCR duplicates were removed with PICARD (version 2.22.8; Broad Institute). SNPs were called with GATK HaplotypeCaller (Poplin et al., 2017) following GATK best practices (Van der Auwera et al., 2013; Van der Auwera and O’Connor, 2020), here with ploidy = 1, resulting in .vcf files for all raw and GATK-filtered variants. The filtered .vcf was then converted to .nexus format with vcf2phylip (Ortiz, 2019), keeping only sites represented in at least 12 accessions (File S2). Phylogenetic analysis was conducted as above, with the exception that the GHOST heterotachy model was not used (as variation below the species level should not be expected to show strong patterns of heterotachy). Instead, the best-fit model was selected from the entire dataset using ModelFinder (Kalyaanamoorthy et al., 2017) under the Bayesian Information Criterion (BIC).

The annotated mitogenome sequence for *M. vimineum* was deposited in NCBI GenBank under accession OQ360108. Raw read data used to build the genome (PacBio, RNAseq), for phylogenomics, and for SNP analysis (DNAseq) was deposited in the NCBI Sequence Read Archive under BioProject PRJNA769079. Supplementary files are available at https://doi.org/10.5281/zenodo.7618370.

## Results

### Organellar genome sequencing and assembly

The final, polished assembly was 478,010 bp in length (Fig. 1A), with overall GC content at 43.7% (41.3% for protein-coding sequences, 53.0% for rRNA genes, and 51.1% for tRNA genes). The initial assembly resulted in six contigs, three of which comprised the plastid genome (large and small single copy regions, inverted repeat), and three of which comprised the mitogenome. The latter were assembled into a single contig based on overlapping ends with the Geneious *de novo* assembler. Despite attempts to circularize the genome with Circlator, a single, “master circle” model could not be constructed. The final genome assembly contained two large, inverted repeats and a single, large, direct repeat: IR1 (28,247 bp), IR2 (2,380 bp), and DR1 (6,462 bp), respectively. Potential secondary structures of the genome model are depicted in Fig. 1B and C. One possible secondary structure (Fig. 1B) consists of large and small single copy regions (which contain copies of IR2 and DR1) and a large IR(1). Another possible structure, considering both large IR sequences, consists of three single copy regions, separated by the two IRs (Fig. 1C). Mean coverage depths of the three regions from Fig. 1B are: 31x (PacBio) and 42x (Illumina) for IR1; and 23x (PacBio) and 21x (Illumina) for the both “single copy” regions.

**Figure 1.**
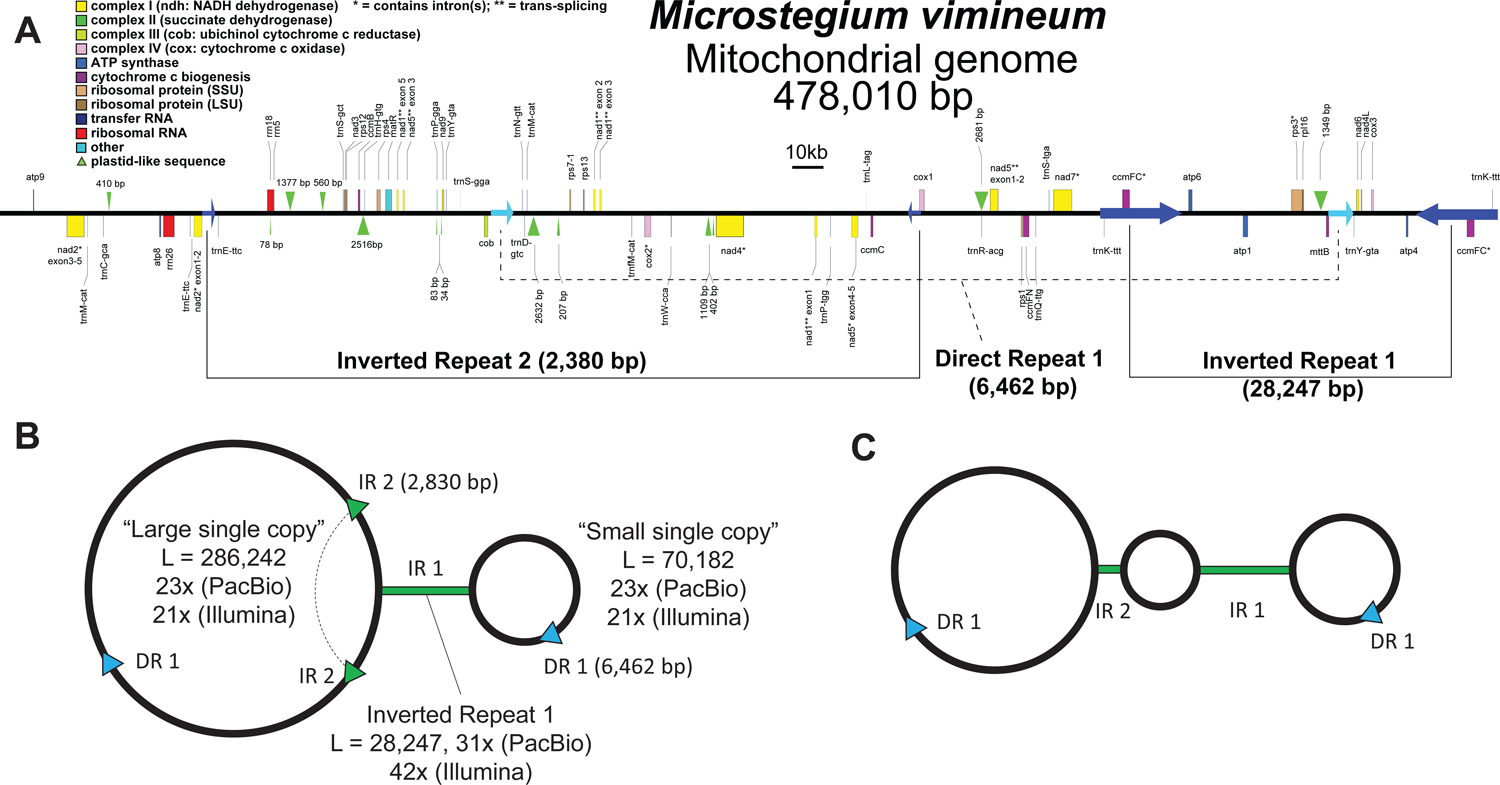
**A**. Linear map of the *Microstegium vimineum* mitogenome assembly and annotation. Scale = 10 kilobases (kb). **B**. Proposed mitogenome secondary structural model emphasizing a single large inverted repeat. **C**. Proposed model emphasizing two large inverted repeats. IR1, IR2 = Inverted Repeat 1 and Inverted Repeat 2 (respectively); DR1 = large, direct repeat 1; L = length of each region; numbers preceding ‘×’ = mean coverage depth of each region in B. Note: the genome in B and C is represented as a looped or circular structure, but the genome could not be circularized with long read data.

### Analyses of repetitive DNA, structural variation, plastid-like DNA, RNA editing, transposable elements, and foreign-acquired sequences

The genome assembly contains 32 protein-coding genes (CDS), three ribosomal RNA genes (*rrn*), and 27 transfer RNA genes (*trn*). In addition to the large repeat regions above, the mitogenome of *M. vimineum* contains numerous smaller repeats (< 1,000 bp). These include direct repeats (880 bp, 262 bp, 164 bp, and 109 bp), inverted repeats (165 bp, 109 bp), and one repeat with three intervals of 154 bp (forward, forward, reverse). The genome contains eight tandem repeat regions: (AC)_6_, (AG)_6_, (AT)_10_, (CT)_6_, and (ACTTT)_5_, and three regions of (AT)_7_. Further, it contains 87 dispersed repeats < 100 bp in length in forward/forward orientation (mean length = 36.3 bp) and 101 in forward/reverse orientation (mean length = 35.0 bp). Comparison of repeat content with relatives of *M. vimineum* within tribe Andropogoneae reveal similar patterns (Fig. 2A): large inverted repeats (i.e. > 5 kb) are present in *M. vimineum* (2), *Chrysopogon zizanioides* (3), and *Coix lacryma*-*jobi* (4). The same is true for large direct repeats, which are present in all species: *Microstegium* (1), *Chrysopogon* (1), *Coix* (2), *Saccharum* (1), and *Sorghum* (2). LastZ alignments revealed seven different structural conformations of the *M. vimineum* mitogenome based on assemblies from eight random subsets of 200,000 PacBio reads (Fig. 2B). Nearly all of the apparent breakpoints were associated with large direct or inverted repeat regions, while one major breakpoint was associated with a small (109 bp) direct repeat.

**Figure 2.**
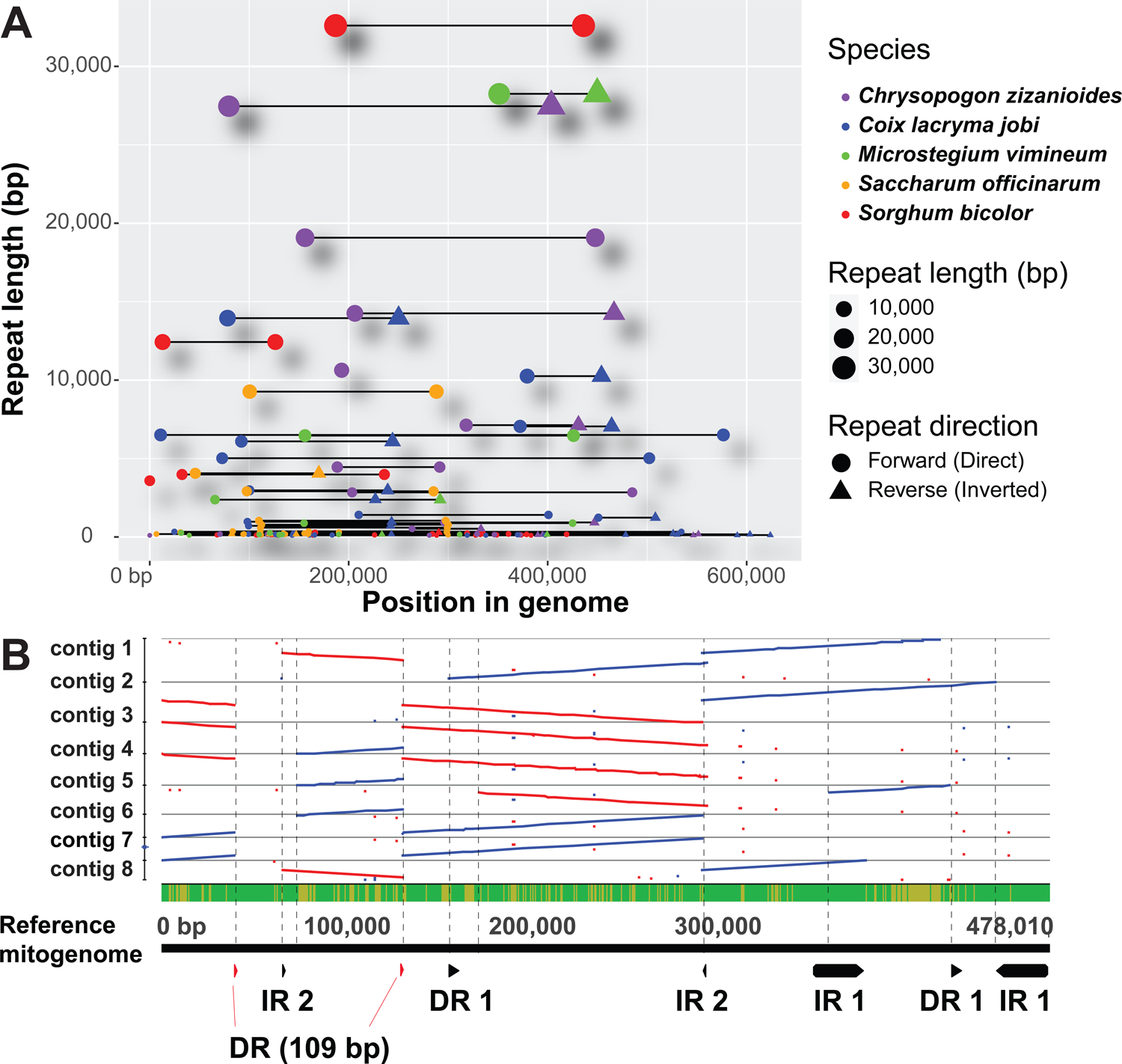
**A**. Repeat distribution in the mitogenomes of five species of grasses within the tribe Andropogoneae. Points represent repeats scaled by size; shapes represent the orientation of each repeat. **B**. Alternative conformations of the mitogenome based on LASTZ alignments of assemblies of eight randomized subsets of PacBio reads, relative to the genome model in Figure 1A. ‘IR’ = inverted repeat, ‘DR’ = direct repeat. Red lines = forward orientation, blue lines = reverse orientation.

Plastid-like sequences comprise 2.4% of the genome. In total, successfully transferred annotations of plastid genes to the mitogenome comprised 43 annotations, 17 of which were CDS (Fig. 3) and the remainder of which were tRNA-like genes. Average percent similarity for CDS was 78.53 (range = 39.7) and for *trn*-like regions was 76.01 (range = 39.2). The three largest annotated plastid-like regions in the *M. vimineum* mitogenome correspond to *atpB*, *psaB*, and *rpoC1*, with percent similarities to their plastomic homologs of 75.3, 73.6, and 70.9, respectively (Fig. 3). Several plastid-like sequences are also found in the mitogenomes of other members of tribe Andropogoneae, and more broadly among grasses, including *atpB, atpE, ndhK, psaB, psbF, rpl14, rpl2, rpl23, rpoC1, rps19,* and *rps2*. All plastid-like mitochondrial regions in *M. vimineum* showed evidence of pseudogenization relative to their plastid-encoded homologs, including high levels of divergence, 5’ or 3’ truncation, internal stop codons, and frameshift insertions or deletions. The only region with an intact reading frame corresponded to *atpE*, having three substitutional differences relative to the plastid copy; two of these were adjacent and resulted in replacements at codons 3 and 4 (L → F and N→H, respectively).

**Figure 3.**
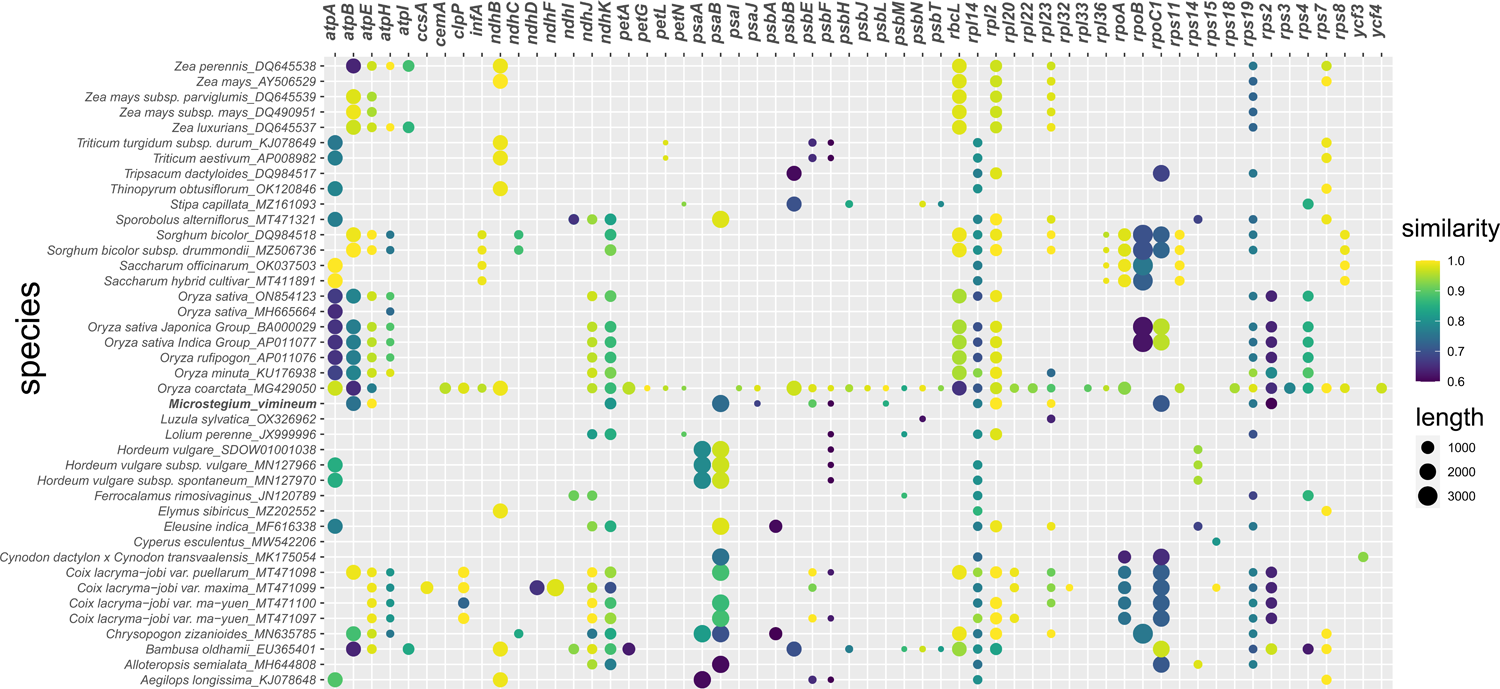
Plastid-like sequences in the mitogenomes of sequenced grass species.

Analyses of gene expression based on RNA-seq data from developing leaf tissue revealed that over half of all expressed mitochondrial transcripts were ATP Synthase (*atp*, TPM range = 33,500.5-329,781.1) (Fig 4A, B), followed by Cytochrome C Oxidase (*cox,* TPM = 30,052.4-67,602.3) and NADH Dehydrogenase (*nad*, 5,668.2-50,473.9) (Fig. 4A, B). Expression was also detected for plastid-like regions, predominantly *ndhK* (TPM = 654,340.2) and *psaJ* (142,073.1) (Fig. 4C). Together, these two regions accounted for >75% of all putatively expressed, plastid-like regions, despite the former having multiple internal stop codons and the latter being truncated at the 3’ end. There was evidence of C→U RNA editing among mitochondrial CDS as well, ranging from 1 site per gene to 18 sites per gene (*viz*. *ccmC*; Fig.4D). The vast majority of RNA editing involved replacement substitutions, with SER→LEU being the most common type (Fig. 4E).

**Figure 4.**
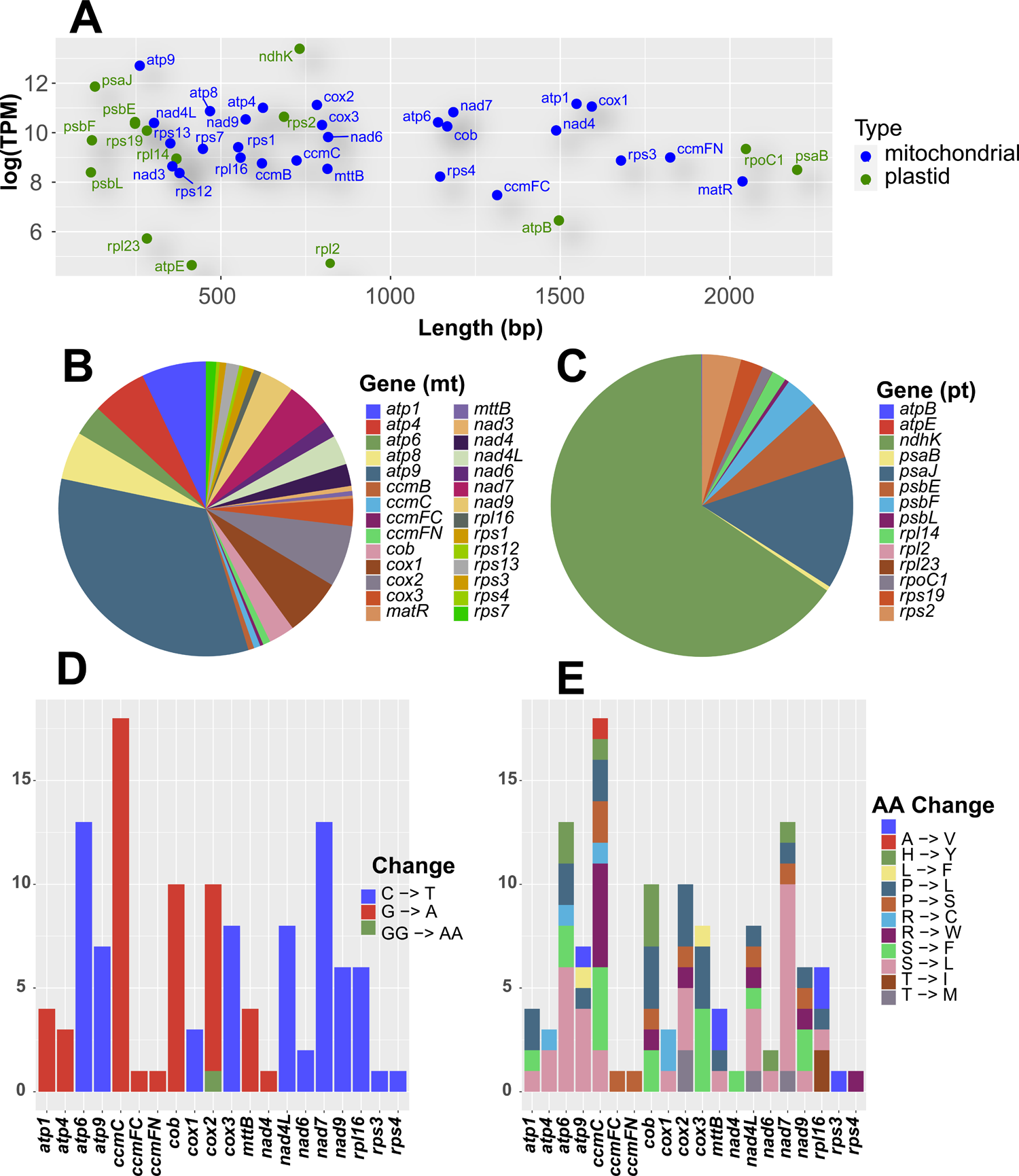
A. Location and relative levels of expression (TPM, transcripts per million) for mitochondrial CDS (coding DNA sequences) and plastid-like sequences. **B**. Relative expression levels of mitochondrial CDS and **C**. plastid-like sequences. **D**. Numbers of putative RNA editing sites for each mitochondrial CDS (C→U on forward strand or G→A on reverse strand). **E**. Predicted amino acid changes at putative RNA editing sites per mitochondrial CDS.

Searches for transposable element-like (TE-like) sequences recovered 29 hits in RepeatMasker, including sequences similar to Class I retrotransposons (n = 14) and Class II DNA transposons (n = 10; Fig. S1). Class I retrotransposon-like sequences belonged to LTR/Copia (n = 4; length range = 102-646 bp), LTR/Gypsy (n = 9; 44-1,313 bp), and LINE/L1 Superfamilies (n = 1; 108 bp). Among Class II DNA transposon-like sequences, nine were similar to DNA/PIF-Harbinger (47-6,881 bp), two of which were > 6 kb in length (Fig. S1). Another Class II-like sequence corresponded to the DNA/CMC-EnSpm superfamily (139 bp). Five TE-like hits were unclassified, ranging from 32-2,437 bp in length. Additional searches with HelitronScanner found three hits for Helitron-like sequences, of 571, 14,875, and 5,477 bp. The first was identified in the spacer region between *rps12* and *ccmB*, the second between *nad4* and *nad1* intron 3, and the third overlapping with *cox1*. MiteFinderII found three hits of MITE-like sequences with length of 246, 400, and 285 bp. The first was identified between *trnS-GGA* and *rps7*, the second between *trnP-TGG* and *nad5* (exon 5), and the third between *ccmFC* and *trnK-TTT* (which is duplicated within the largest inverted repeat region). Taken together, 9.05% of the *M. vimineum* mitogenome is composed of TE-like sequences.

Analysis of the mitogenome assembly with Kraken2 supports a genome which is free from foreign sequences and contamination (Fig. S2). Scanning of k-mers comprising 100 bp segments of the mitogenome against those found in genomes representing plants, bacteria, archaea, viruses, fungi, UniVec contaminants, and the human genome resulted in classification of 77.52% of k-mers, of which 77.45% were classified as characteristic of k-mers optimized to the node representing the hypothesized ancestor of Viridiplantae, 74.44% of Liliopsida, and 72.3% as Poaceae. The remaining unclassified k-mers either represent our incomplete knowledge of mitogenomic diversity or the presence of sequences unique to the mitogenome of this species.

### Phylogenomic analyses using mitochondrial CDS

Analysis of 7,019 aligned positions across 28 protein-coding mitochondrial genes (total gaps/missing data content = 7.2 %, total parsimony-informative characters = 1,955) under the GHOST heterotachy model yielded a tree topology with generally high bootstrap support values (lnL = −31,900.1643, BIC score = 65,013.6522, total branch/model free parameters = 137; Fig. 5A). Among the families of order Poales, mitochondrial data placed Typhaceae as sister to Bromeliaceae + the remainder of the order (BS = 99). In the latter clade, Xyridaceae were sister to a clade composed of ((Mayacaceae, (Thurniaceae, Cyperaceae, Juncaceae)), (Joinvilleaceae, Poaceae)); all BS = 100 excluding the sister relationship of (*Mayaca, Xyris*) (BS = 47) and Mayacaceae as sister of (Thurniaceae, Cyperaceae, Juncaceae) (BS = 77). Within Poaceae, *Puelia* (Puelioideae) was supported as sister to the remaining taxa (BS = 100), followed by representatives of tribes Bambusoideae (*Bambusa*, *Ferrocalamus*) and tribe Pooideae (e.g. *Lolium*, *Triticum*, *Thinopyrum*, *Hordeum*), but with a lack of support for the latter subfamilies as sister to one another (BS = 48). Following this, Oryzoideae (*Oryza*) (BS = 100) was placed as sister to Chloridoideae (*Eleusine*, BS = 100), and Panicoideae (BS = 100). Within Panicoideae, *Alloteropsis* was sister to a clade comprising members of the subtribe Andropogoneae (BS = 99). Within Andropogoneae, (*Tripsacum*, *Zea*) were sister to a clade composed of *Sorghum*, *Microstegium*, *Saccharum*, *Chrysopogon*, and *Coix* (BS = 100), with BS = 92 for the latter. However, support was generally low within this clade. *Microstegium nudum* was placed as sister to two accessions of *Sorghum*, but with no support (BS = 52). Sister to this clade is a clade of (*Coix, Microstegium*), but again, with no support (BS = 46). Among the remaining accessions of *Microstegium, M. faurei* was sister to the rest but with no support (BS = 52), while *M. vimineum* was placed as sister to *M. japonicum* and an unknown accession of *Microstegium* from Yunnan, China (BS = 100); the latter specimen was 100% identical to *M. japonicum*.

**Figure 5.**
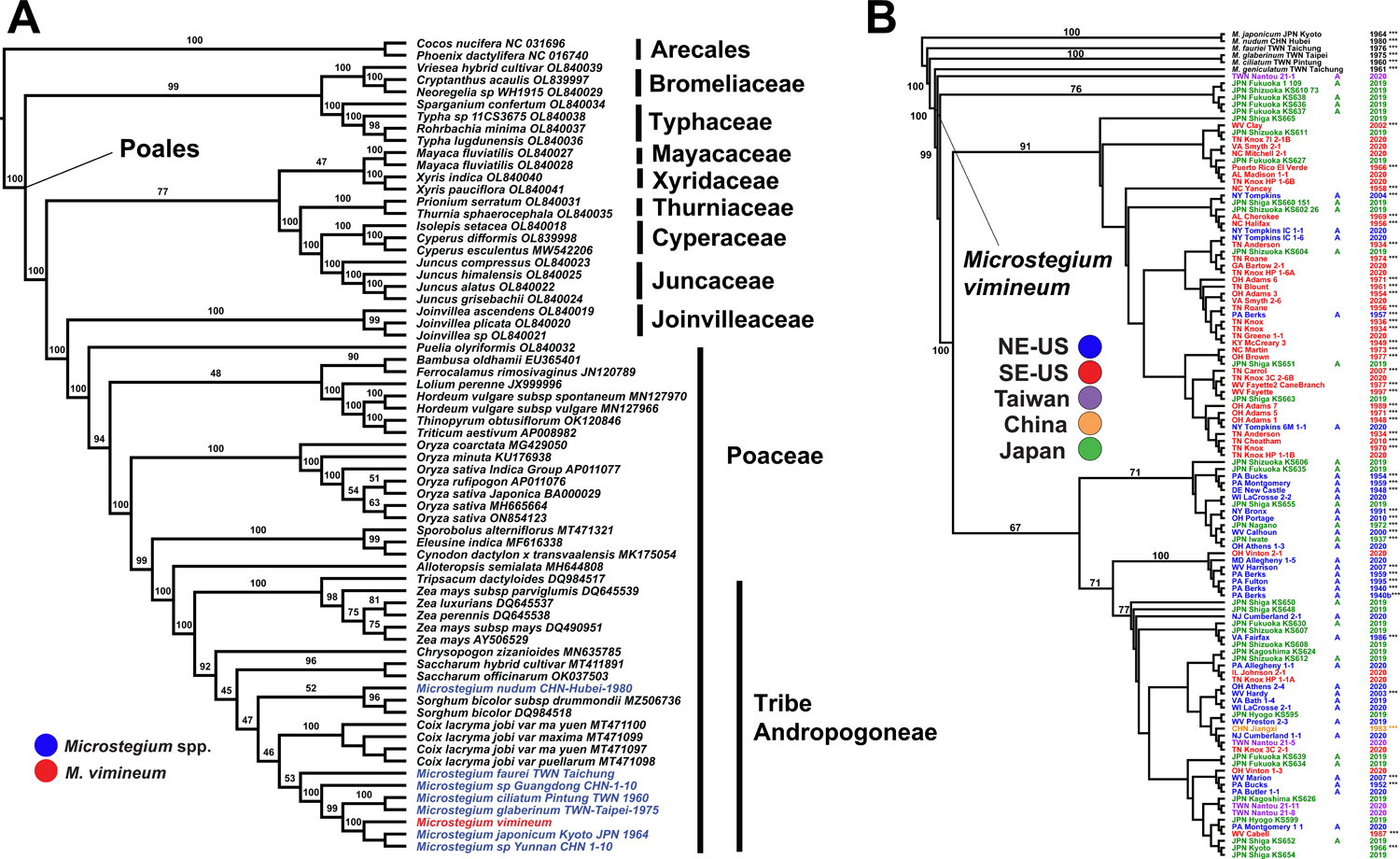
**A**. Maximum likelihood phylogenetic tree based on a 7,019 bp alignment of mitochondrial CDS under the GHOST heterotachy model. Numbers above branches indicate bootstrap support. *Microstegium* species are in blue font, with *M. vimineum* in red. **B**. Maximum likelihood phylogenetic tree among mitochondrial haplotypes of *M. vimineum*. Colors indicate regions from which samples were collected (green = Japan, orange = China, purple = Taiwan, red = the southeastern US, blue = the northeastern US). ‘A’ indicates the presence of awned florets, and a lack thereof indicates a lack of awned florets. The numbers on the right list the year each sample was collected; all samples collected prior to 2019 came from herbarium specimens, indicated by ‘***’.

### Assessment of relationships among mitochondrial haplotypes of M. vimineum

Analysis of genome skim datasets for 118 accessions from the US (invasive) and Asia (native) yielded 3,913 mitochondrial variants. Phylogenetic analysis of the data in IQtree2 yielded a tree with two principal clades corresponding to samples from the invasive range (best-fit model = SYM+ASC+R5, lnL = −47,277.03, BIC score = 96,588.99, total branch/model free parameters = 246; Fig. 5B). These two clades were sister to a clade of haplotypes from Japan (from Fukuoka, Shiga, and Shizuoka), collectively sister to a single haplotype from Nantou, Taiwan. The first clade containing accessions from the invasive range was primarily composed of individuals from the southeastern USA that lack awns (BS = 91). Interspersed among these invasive-range accessions were several accessions from Japan. Bootstrap values among individual haplotypes within this clade were generally low. The second clade was composed of primarily awned forms from the northeastern US, but this clade as a whole received weak support (BS = 67). Invasive haplotypes in this clade were interspersed among those from Japan and Taiwan, with a single haplotype from China; likewise, support values were generally low within this clade. There were exceptions, however: four haplotypes from Tompkins County, New York, USA grouped with the predominantly “southern” clade (predominantly southern US awnless accessions), whereas six haplotypes from eastern Tennessee, southern West Virginia, southern Ohio, and southern Illinois, USA grouped with the predominantly “northern” clade.

## Discussion

### Organellar genome sequencing and assembly

We sequenced and analyzed the 478,010 bp mitochondrial genome of the invasive *M. vimineum*, revealing a genome typical of previously sequenced grasses. Grass mitogenomes represented in NCBI GenBank range from 294 to 740 kb (last accessed January, 2023), thus *M. vimineum* has a somewhat average genome size with gene content typical of other grasses. Overall, gene space occupies 13.2% of the genome, followed by TE-like sequence (9.05%) and plastid-like sequence (2.4%), leaving 75.3% as unknown.

### Analyses of repetitive DNA, structural variation, plastid-like DNA, RNA editing, transposable elements, and foreign-acquired sequences

As observed in other mitogenomes, both large (i.e > 1,000 bp) and small direct and indirect repeats are present in *M. vimineum* (Figs. 1, 2). Further, these repeats are associated with putative isomeric variants, which argues against the existence of a “master circle” (Figs. 1, 2; Sloan, 2013). In fact, the genome could not be circularized with PacBio or Illumina reads, casting further doubt on the existence of a single circular structure. Insertions of plastid-like DNA regions, many of which are divergent from their homologs in the plastid genome of *M. vimineum*, suggests that many of these regions may be considered “ancient” transfers, while some others may have either occurred more recently or are the result of “copy correction” via gene conversion (Fig. 3A; Sloan and Wu, 2014). The total extent of plastid DNA content detected in the mitogenome species is not extreme (2.4% compared to > 10% in the palm *Phoenix dactylifera*; Fang et al., 2012), but is similar to that in other grasses (e.g. Clifton et al., 2004). The apparent expression of some of these regions presents a conundrum, as our evidence suggests these are non-functional, lacking intact open reading frames. One possible explanation would be that these plastid-like regions lie within expressed cistrons, and thus are transcribed but potentially spliced out or their RNAs modified post-transcription (Cardi et al., 2012). Previous research has shown that most of the mitogenome can be transcribed, and that extensive post-transcriptional modification produces the mature transcripts (Holec et al., 2006; Ruwe et al., 2016). RNA editing was also observed within CDS of *M. vimineum*, a common feature of both organellar genomes in plants; this process is likely essential for proper gene expression and further may preserve the integrity of secondary structure in organellar genomes (e.g. Maier et al., 1996). It should be noted that only a single tissue type (young developing leaf tissue) from a single individual was included here, and thus the need remains for gene expression studies across tissue types, developmental stages, and environmental conditions to explore the transcriptional landscape in this invasive species.

Integration of nuclear-derived TE-like sequences provide a partial explanation for plant mitochondrial genome size expansion (Marienfeld et al. 1999, Mower et al. 2012, Zhao et al. 2018). Previous research on *Arabidopsis thaliana*, *Citrullus lanatus*, *Cucurbita pepo*, *Lingustrum quihoui* and, *Elymus sibiricus* have reported ∼1-6% of nuclear-derived TE-like sequences in their respective mitogenomes (Knoop et al. 1996, Alverson et al, 2010, Yu et al. 2020, Xiong et al. 2022). Although 9.05% of the mitogenome of *M*. *vimineum* is occupied by similar TE-like sequences (Fig. S1), the majority of them are fragmented copies. These results indicate frequent and independent DNA transfers from nuclear to mitochondrial genome, that the fragmented copies could have either been generated from former complete sequences that later became degraded, they originated from incomplete transposition events, or they were scrambled by intramolecular recombination which is frequent in plant mitogenomes (Knoop et al. 1996, Notsu et al. 2002). Regardless, the landscape of TE-like sequences in plant mitogenomes is not well explored.

An absence of sequence from distantly related plant lineages, or other lineages in general, suggests that the mitogenome of *M*. *vimineum* is free of foreign sequence, foreign sequence is too recombined to identify, or k-mers present are highly unique and are not contained in the genomes included in our analyses (Fig. S2). We find the latter two explanations unlikely, given the broad range of lineages represented in our Kraken2 databases. Additionally, none of the k-mers from our assembly were classified when searched against the Eukaryotic Pathogen, Vector and Host informatics Resource Database. Taken together, these results characterize a mitogenomic assembly which is free of confounding artifactual contamination resulting from interactions in the lab or during necessary bioinformatic components of our work.

### Phylogenomic analyses using mitochondrial CDS

*Microstegium* species, including *M. vimineum*, are clearly placed within the grass tribe Andropogoneae based on mitochondrial data (Fig. 5A). Our analysis of mitochondrial coding regions suggests a close relationship among most of the *Microstegium* species sampled here, with the exclusion of *M. nudum*, and possibly *M. faurei*; the latter was placed as sister of *M. ciliatum*, *M. glaberrimum*, *M. japonicum*, and *M. vimineum*, but with no support. This is in contrast to other studies based on plastid DNA in which species of *Microstegium* occupy different clades within the Andropogoneae, though the level of *Microstegium* spp. sampling in those studies, and the lack of available mitogenomes across Andropogoneae in the current study, are insufficient for confident placement of the different species. Lloyd Evans et al. (2019) placed *M. vimineum* with moderate support as sister to *Polytrias* and two species of *Sorghastrum*, all of which are sister to a clade composed of *Miscanthus* and *Saccharum* spp. based on five low-copy nuclear genes. In that study, *M. vimineum* is estimated to have diverged from a common ancestor with *Polytrias* and *Sorghastrum* between 7 and 10.5 million years ago, but sampling only included *M. vimineum* from the genus *Microstegium*. Data from complete plastid genomes placed *M. vimineum* as sister to two genera: *Kerriochloa*, with a single species *K. siamensis* (Thailand, Vietnam); and *Sehima*, comprising five species from Africa, Asia, and Australia (Welker et al., 2020). But again, *M. vimineum* was the only representative of *Microstegium* sampled.

Chen et al. (2009; 2012) conducted a phylogenetic analysis and taxonomic treatment of *Microstegium* based on nuclear ITS sequencing and morphology. Our findings of *M. nudum* as sister of *Sorghum*, with other species of *Microstegium* occupying a different clade (more closely allied with *Coix*) are generally in agreement with these previous studies, but with some key differences. First, Chen et al. (2012) identify two clades, a ‘*M. nudum*’ clade (*japonicum*, *nudum*, *somae*), and a ‘*M. vimineum*’ clade (*ciliatum, faurei, geniculatum, vimineum*). In our phylogenetic analysis, only *M. nudum* grouped outside the main clade of *Microstegium*, but support overall for the latter is weak (Fig. 5A). In our analysis, *M. vimineum* was strongly supported as sister to *M. japonicum* and an unknown species of *Microstegium* (BS = 100), whereas the analysis of Chen et al. (2012) placed *M. vimineum* as sister to *M. ciliatum*, but with low support (BS = 69). While the taxon sampling of mitogenomes in the current study is not comprehensive, it does represent the largest amount of data analyzed to date on the taxonomic status of the genus.

Based on our analysis and previous studies, it is indeed possible that *Microstegium* is non-monophyletic, perhaps reflecting a complex, reticulate history of allopolyploidy that is broadly observed among the Andropogoneae (e.g. Estep et al., 2014; Hawkins et al., 2015; Arthan et al., 2017; Ramachandran et al., 2021). *Microstegium vimineum* is a known polyploid, with 2N = 20 chromosomes, twice that of the “base” number of 2N = 10 in Andropogoneae (Watson and Dallwitz, 1992). Further, analysis of the recently published chromosome-level nuclear genome of *M. vimineum* revealed strong evidence of a paleopolyploidy event, with about 1/3 of all nuclear genes present as duplicate copies (Ramachandran et al., 2021). Further, comparative analysis of terminal repeats of transposable elements throughout the nuclear genome, calibrated to a grass-specific TE divergence rate (Ma and Bennetzen, 2004), revealed a burst of TE activity roughly 1-2 million years ago, possibly coinciding with “genomic shock” associated with a polyploidy event (Ramachandran et al., 2021). This warrants further study with dense taxon sampling across the tribe, including multiple species and accessions of *Microstegium*, and employing genome-wide plastid, mitochondrial, and nuclear markers to test hypotheses of allopolyploid origins within the currently circumscribed *Microstegium* and other genera.

### Assessment of relationships among mitochondrial haplotypes of M. vimineum

Patterns of mitogenomic SNP variation within *M. vimineum* reveal a complex invasion history (Fig. 5B). The finding of a predominantly “northern awned” clade and a “southern awnless” clade mirrors that based on nuclear SNP data (Barrett et al., 2022) and plastid data (Corbett C. W. et al., unpublished data). Further, there is evidence of multiple invasions and subsequent establishments from the native range, with a likely initial, successful “awnless” invasion in the southeastern US and at least one more successful invasion in the northeastern US of the awned form, likely in eastern Pennsylvania. Because this species was used as packing for shipments from Asia, it is plausible for multiple invasions to have occurred, perhaps bearing higher than expected genetic diversity (i.e. contrasted with expectations of a severe genetic bottleneck upon a single invasion) if seeds from multiple plants continually became established (e.g. Sakai et al., 2001; Kolbe et al., 2004; Frankham, 2005; Dlugosch and Parker, 2007; Sutherland et al., 2021). Further, there is evidence that each putative invasion and subsequent spread led to long-distance dispersal within the invasive range over the last century, with “southern” mitotypes present as far north as central-western New York State (i.e. Tompkins County, New York), and “northern” mitotypes present as far south as eastern Tennessee (Fig. 5B). Indeed, samples collected in 2020 from the original site where stiltgrass was collected 101 years before (Knox County, Tennessee) revealed a mix of northern and southern mitotypes, suggesting that this species has been dispersed extensively over the past few decades via anthropogenic activity. This is significant, as such long-distance dispersal may lead to rapid admixture of previously separated genotypes from the native range, allowing the genomic potential for rapid adaptation to local conditions (Verhoeven et al., 2010; Rius and Darling, 2014), and thus presenting a mechanism for increased invasive potential over time (Dlugosch and Parker, 2008; Keller and Taylor, 2010; Dlugosch et al., 2015; Sutherland et al., 2021).

There is evidence that *M. vimineum* has experienced rapid adaptation after becoming established in the invasive range, in terms of flowering phenology across a latitudinal gradient, and growth/reproductive advantages of invasive populations compared with those in the native range, in line with the Evolution of Increased Competitive Ability hypothesis (Flory et al., 2011; Novy et al., 2013; Huebner et al., 2022). Barrett et al (2022) suggested that this may further extend to selection in the invasive range for different awn phenotypes. In the eastern US, there is a strong latitudinal pattern of awnless forms in the South, long-awned forms in the North, and intermediate- or short-awned forms at mid-latitudes. A similar but relatively weaker pattern was observed in Asia, with both awned and awnless phenotypes intermixed at low and mid-latitudes, but a predominance of awned forms at higher latitudes. Awns are hypothesized to aid in microsite dispersal and burial via hygroscopic movement, effectively drilling the seed-containing floret into the seed bank (Cavanagh et al., 2020). Awns are expected to play a role in seedling burial and increased survival from frequent and intense soil freezing events at higher latitudes.

Our mitochondrial SNP analysis (Fig. 5B), and previous analysis of nuclear SNP variation (Barrett et al., 2022), support a scenario consistent with intensified, post-invasion selection for awn phenotypes in the eastern US, favoring awnless forms at lower latitudes and awned forms at higher latitudes. Habitat filtering may have also played a role (Weiher and Keddy, 1995), by selecting which phenotypes were successful in their initial invasions, with a higher likelihood of successful invasion hypothesized in the South by awnless forms, and by awned forms in the North. We are currently conducting burial, germination, and seed survival experiments to test hypotheses on selection for awn phenotypes in the invasive range.

Genome sequencing efforts in invasive plant species are in their infancy, but hold great potential for the identification, phylogenetic placement, evolutionary ecology, and management of these species. High-quality genome sequences and annotations provide much needed baseline data, enabling subsequent studies of invasion routes, invasion history, and other diverse applications in invasion biology. Here we have sequenced a reference mitogenome for *M. vimineum*, one of the most damaging invasive plants in North America, to aid in such future studies. While characterizing genome structure and content, we also corroborated recent studies on the complex invasion history and spatiotemporal patterns of mitogenomic variation in the native and invasive ranges. Most importantly, such genomic resources will aid in efforts to predict ongoing patterns of spread within this species, responses to climate change, and possibly help predict future threats from other invasive species, allowing genomically informed forecasting.

## Author contributions

CFB conceived the study, analyzed data, and led the writing of the manuscript. KS, W-BY, and C-HC provided Asian samples, and contributed to drafts of the manuscript. DR and BTS analyzed data and helped draft the manuscript. CDH collected seed, maintained plants in a controlled environment (growth chamber), and helped draft the manuscript. CWC generated data, assisted with data analyses, and helped draft the manuscript. All authors have reviewed and approved the final manuscript.

## Acknowledgments

This work was supported by the US National Science Foundation (award OIA-1920858), the West Virginia University Department of Biology, and the USDA Forest Service Northern Research Station. The authors thank Mark Daly, Jasmine Haimovitz, Joanna Gallagher, and Jordan Zhang at Dovetail Genomics, LLC (Cantata Bio, LLC) for expert assistance with long-read sequencing, and GeneWiz, Inc. for RNA sequencing. We thank J. Beck, M. Latvis, N. Kooyers, M. McKain, E. Sigel, and B. Sutherland for feedback and discussion. We thank the following collaborators for providing contemporary field-collected material: M. McKain, G. Matlack, G. Moore, S. Kuebbing, B. Molano-Flores, A. Kennedy, J. Fagan, N. Koenig, P. Crim, G. Scott, B. Foster, M. Heberling, A. Bowe, P. Wolf, K. Willard, and J. McNeal. For access to herbarium collections we thank: Tiana Rehman (BRIT), Bonnie Isaac (CM), Mason Heberling (CM), John Freudenstein (OS), Anna Statler (BH), Tanya Livshultz (PH), Meghann Toner (US), Lauren Boyle (MO), Margaret Oliver (TENN), and Donna Ford-Werntz (WVA). For assistance with genomic sequencing, we thank R. Percifield, D. Primerano, and J. Fan. For collections in Japan, we thank C. Hara, S. Mori, M. Sato, T. Shimizu, and K. Tanaka. We thank the WVU Genomics Core Facility for support provided to help make this publication possible and for CTSI Grant no. U54 GM104942, which in turn provides financial support to the WVU Core Facility. We further acknowledge WV-INBRE (P20GM103434), a COBRE ACCORD grant (1P20GM121299), and a West Virginia Clinical and Translational Science Institute (WV-CTSI) grant (2U54GM104942) in supporting the Marshall University Genomics Core (Research Citation: Marshall University Genomics Core Facility, RRID:SCR_018885).

## Appendix A1

Voucher information: Species, Accession, US County (if applicable), Region/State, Country, Herbarium Code, Collector, Collection Number. Herbarium Codes: BH (Bailey Hortorium Herbarium), BRIT (Botanical Research Institute of Texas), CM (Carnegie Museum of Natural History), MO (Missouri Botanical Garden), OS (Ohio State University Herbarium), PH (Academy of Natural Sciences), TENN (University of Tennessee Herbarium), WVA (West Virginia University Herbarium).

*Microstegium nudum* (Trin.) A. Camus, M-nudum-CHN-Hubei-1980-83-S44, n/a, Hubei, China, 1980, CM, Bartholomew et al., s.n.; *Microstegium vimineum* (Trin.) A. Camus, CHN-Jiangxi-1983-12-S7, n/a, Jiangxi, China, 1983, CM, Yao, 8711; *Microstegium vimineum* (Trin.) A. Camus, 111-JPN-Fukuoka-KS630-S9, n/a, Fukuoka, Japan, 2019, WVA, Suetsugu, 630; *Microstegium vimineum* (Trin.) A. Camus, 115-JPN-Fukuoka-KS636-S11, n/a, Fukuoka, Japan, 2019, WVA, Suetsugu, 636; *Microstegium vimineum* (Trin.) A. Camus, 116-JPN-Fukuoka-KS637-S12, n/a, Fukuoka, Japan, 2019, WVA, Suetsugu, 637; *Microstegium vimineum* (Trin.) A. Camus, 117-JPN-Fukuoka-KS638-S13, n/a, Fukuoka, Japan, 2019, WVA, Suetsugu, 638; *Microstegium vimineum* (Trin.) A. Camus, 118-JPN-Fukuoka-KS639-S14, n/a, Fukuoka, Japan, 2019, WVA, Suetsugu, 639; *Microstegium vimineum* (Trin.) A. Camus, JPN-Fukuoka-1-109-S58, n/a, Fukuoka, Japan, 2019, WVA, Suetsugu, 633; *Microstegium vimineum* (Trin.) A. Camus, JPN-Fukuoka-KS627-30-S17, n/a, Fukuoka, Japan, 2019, WVA, Suetsugu, 627; *Microstegium vimineum* (Trin.) A. Camus, JPN-Fukuoka-KS634-31-S18, n/a, Fukuoka, Japan, 2019, WVA, Suetsugu, 634; *Microstegium vimineum* (Trin.) A. Camus, JPN-Fukuoka-KS635-32-S19, n/a, Fukuoka, Japan, 2019, WVA, Suetsugu, 635; *Microstegium vimineum* (Trin.) A. Camus, JPN-Shizuoka-KS604-67-S28, n/a, Fukuoka, Japan, 2019, WVA, Suetsugu, 604; *Microstegium vimineum* (Trin.) A. Camus, JPN-Hyogo-KS595-24-S11, n/a, Hyogo, Japan, 2019, WVA, Suetsugu, 595; *Microstegium vimineum* (Trin.) A. Camus, JPN-Hyogo-KS599-25-S12, n/a, Hyogo, Japan, 2019, WVA, Suetsugu, 599; *Microstegium vimineum* (Trin.) A. Camus, JPN-Kagoshima-KS624-28-S15, n/a, Kagoshima, Japan, 2019, WVA, Suetsugu, 624; *Microstegium vimineum* (Trin.) A. Camus, JPN-Kagoshima-KS626-29-S16, n/a, Kagoshima, Japan, 2019, WVA, Suetsugu, 626; *Microstegium japonicum* (Miq.) Koidz., M-japonicum-JPN-Kyoto-1964-84-S45, n/a, Kansai, Japan, 1964, CM, Murata, 19181; *Microstegium vimineum* (Trin.) A. Camus, JPN-Kyoto-1966-11-S6, n/a, Kansai, Japan, 1966, CM, Murata, G., 19905; *Microstegium vimineum* (Trin.) A. Camus, 92-JPN-Nagano-1972-S4, n/a, Nagano, Japan, 1972, CM, Shimizu, T., 24216; *Microstegium vimineum* (Trin.) A. Camus, 120-JPN-Shiga-KS648-S16, n/a, Shiga, Japan, 2019, WVA, Suetsugu, 648; *Microstegium vimineum* (Trin.) A. Camus, JPN-Shiga-KS650-122-S61, n/a, Shiga, Japan, 2019, WVA, Suetsugu, 650; *Microstegium vimineum* (Trin.) A. Camus, JPN-Shiga-KS651-123-S62, n/a, Shiga, Japan, 2019, WVA, Suetsugu, 651; *Microstegium vimineum* (Trin.) A. Camus, JPN-Shiga-KS652-124-S63, n/a, Shiga, Japan, 2019, WVA, Suetsugu, 652; *Microstegium vimineum* (Trin.) A. Camus, JPN-Shiga-KS654-126-S65, n/a, Shiga, Japan, 2019, WVA, Suetsugu, 654; *Microstegium vimineum* (Trin.) A. Camus, JPN-Shiga-KS655-127-S66, n/a, Shiga, Japan, 2019, WVA, Suetsugu, 655; *Microstegium vimineum* (Trin.) A. Camus, JPN-Shiga-KS660-151-S76, n/a, Shiga, Japan, 2019, WVA, Suetsugu, 660; *Microstegium vimineum* (Trin.) A. Camus, JPN-Shiga-KS663-154-S79, n/a, Shiga, Japan, 2019, WVA, Suetsugu, 663; *Microstegium vimineum* (Trin.) A. Camus, JPN-Shiga-KS665-156-S81, n/a, Shiga, Japan, 2019, WVA, Suetsugu, 665; *Microstegium vimineum* (Trin.) A. Camus, JPN-Shizuoka-KS602-26-S13, n/a, Shizuoka, Japan, 2019, WVA, Suetsugu, 602; *Microstegium vimineum* (Trin.) A. Camus, JPN-Shizuoka-KS606-69-S30, n/a, Shizuoka, Japan, 2019, WVA, Suetsugu, 606; *Microstegium vimineum* (Trin.) A. Camus, JPN-Shizuoka-KS607-70-S31, n/a, Shizuoka, Japan, 2019, WVA, Suetsugu, 607; *Microstegium vimineum* (Trin.) A. Camus, JPN-Shizuoka-KS608-71-S32, n/a, Shizuoka, Japan, 2019, WVA, Suetsugu, 608; *Microstegium vimineum* (Trin.) A. Camus, JPN-Shizuoka-KS610-73-S34, n/a, Shizuoka, Japan, 2019, WVA, Suetsugu, 610; *Microstegium vimineum* (Trin.) A. Camus, JPN-Shizuoka-KS611-74-S35, n/a, Shizuoka, Japan, 2019, WVA, Suetsugu, 611; *Microstegium vimineum* (Trin.) A. Camus, JPN-Shizuoka-KS612-27-S14, n/a, Shizuoka, Japan, 2019, WVA, Suetsugu, 612; *Microstegium vimineum* (Trin.) A. Camus, JPN-Iwate-1937-82-S43, n/a, Tohoku, Japan, 1937, CM, Iwabuchi, H., 5516; *Microstegium vimineum* (Trin.) A. Camus, TWN-N-21-11-S38, n/a, Nantou, Taiwan, 2020, WVA, Chen, TWN-N-21-11; *Microstegium vimineum* (Trin.) A. Camus, TWN-N-21-2-S29, n/a, Nantou, Taiwan, 2020, WVA, Chen, TWN-N-21-2; *Microstegium vimineum* (Trin.) A. Camus, TWN-N-21-5-S32, n/a, Nantou, Taiwan, 2020, WVA, Chen, TWN-N-21-5; *Microstegium vimineum* (Trin.) A. Camus, TWN-N-21-8-S35, n/a, Nantou, Taiwan, 2020, WVA, Chen, TWN-N-21-8; *Microstegium ciliatum* (Trin.) A. Camus, M-ciliatum-TWN-Pintung-1960-88-S49, n/a, Pintung, Taiwan, 2020, CM, Hsu, 1073; *Microstegium fauriei* (Hayata) Honda), M-fauriei-TWN-Taichung-1976-87-S48, n/a, Taichung, Taiwan, 2020, CM, Kuo, 7116; *Microstegium geniculatum* (Hayata) Honda, M-geniculatum-TWN-Taichung-1961-86-S47, n/a, Taichung, Taiwan, 2020, CM, Feung, 4439; *Microstegium glaberrimum* (Honda) Koidz., M-glaberinum-TWN-Taipei-1975-85-S46, n/a, Taipei, Taiwan, 2020, CM, Kuo, 6424; *Microstegium vimineum* (Trin.) A. Camus, AL-Cherokee-1969-S67, Cherokee, Alabama, USA, 2020, TENN, Kral, 37767; *Microstegium vimineum* (Trin.) A. Camus, AL-MAD-1-1-S51, Madison, Alabama, USA, 2020, WVA, Wolf, AL-MAD-1-1; *Microstegium vimineum* (Trin.) A. Camus, DE-New-Castle-1948-138-S68, New Castle, Delaware, USA, 1948, CM, Long, B., 68410; *Microstegium vimineum* (Trin.) A. Camus, GA-BAR-2-1-S55, Bartow, Georgia, USA, 2020, WVA, McNeal, GA-BAR-2-1; *Microstegium vimineum* (Trin.) A. Camus, IL-JOHN-2-1-S15, Johnson, Illinois, USA, 2020, WVA, Molano-Flores, IL-JOHN-2-1; *Microstegium vimineum* (Trin.) A. Camus, KY-McCreary3-1949-23-S10, McCreary, Kentucky, USA, 1949, MO, Reed, 16339; *Microstegium vimineum* (Trin.) A. Camus, 215-MD-GR-1-5-S15, Alleghany, Maryland, USA, 2019, WVA, Huebner, MD-GR-1-5; *Microstegium vimineum* (Trin.) A. Camus, NJ-CUMB-1-1-S5, Cumberland, New Jersey, USA, 2020, WVA, Moore, NJ-CUMB-1-1; *Microstegium vimineum* (Trin.) A. Camus, NJ-CUMB-2-2-S6, Cumberland, New Jersey, USA, 2020, WVA, Moore, NJ-CUMB-2-2; *Microstegium vimineum* (Trin.) A. Camus, NY-Bronx-1991-140-S69, Bronx, New York, USA, 1991, CM, Nee, M., 41826; *Microstegium vimineum* (Trin.) A. Camus, NY-Thomkins-2004-S86, Tompkins, New York, USA, 2004, BH,; *Microstegium vimineum* (Trin.) A. Camus, NY-TOM-6M-1-1-S20, Tompkins, New York, USA, 2020, WVA, Bowe, NY-TOM-6M-101; *Microstegium vimineum* (Trin.) A. Camus, NY-TOM-IC-1-1-S18, Tompkins, New York, USA, 2020, WVA, Bowe, NY-TOM-IC-1-1; *Microstegium vimineum* (Trin.) A. Camus, NY-TOM-IC-1-6-S19, Tompkins, New York, USA, 2020, WVA, Bowe, NY-TOM-IC-1-6; *Microstegium vimineum* (Trin.) A. Camus, NC-Halifax-1956-50-S21, Halifax, North Carolina, USA, 1956, BRIT, Ahles, 20724; *Microstegium vimineum* (Trin.) A. Camus, NC-Yancey-1958-51-S22, Yancey, North Carolina, USA, 1958, BRIT, Ahles, 50776; *Microstegium vimineum* (Trin.) A. Camus, NC-Martin-1973-76-S37, Martin, North Carolina, USA, 1973, CM, Boufford, D.E., 12249; *Microstegium vimineum* (Trin.) A. Camus, NC-SP-2-1-S57, Mitchell, North Carolina, USA, 2020, WVA, Barrett, NC-SP-2-1; *Microstegium vimineum* (Trin.) A. Camus, OH-Adams-Co1-1948-4-S4, Adams, Ohio, USA, 1948, OS, Barley, F., s.n.; *Microstegium vimineum* (Trin.) A. Camus, OH-Adams-Co3-1954-5-S5, Adams, Ohio, USA, 1954, OS, Barley, F., s.n.; *Microstegium vimineum* (Trin.) A. Camus, OH-Adams-Co5-1971-1-S1, Adams, Ohio, USA, 1971, OS, Barley, F., s.n.; *Microstegium vimineum* (Trin.) A. Camus, OH-Adams-Co6-1971-2-S2, Adams, Ohio, USA, 1971, OS, Barley, F., s.n.; *Microstegium vimineum* (Trin.) A. Camus, 89-Brown-OH-1977-S1, Brown, Ohio, USA, 1977, OS, Cusick, A.W., s.n.; *Microstegium vimineum* (Trin.) A. Camus, OH-Adams-Co7-1989-3-S3, Adams, Ohio, USA, 1989, OS, Barley, F., s.n.; *Microstegium vimineum* (Trin.) A. Camus, 91-Portage-OH-2010-S3, Portange, Ohio, USA, 2010, OS, Gardner, R.L., 6970; *Microstegium vimineum* (Trin.) A. Camus, OH-ATH-1-3-S1, Athens, Ohio, USA, 2020, WVA, Matlack, OH-ATH-1-3; *Microstegium vimineum* (Trin.) A. Camus, OH-ATH-2-4-S2, Athens, Ohio, USA, 2020, WVA, Matlack, OH-ATH-2-4; *Microstegium vimineum* (Trin.) A. Camus, OH-VIN-1-3-S12, Vinton, Ohio, USA, 2020, WVA, Scott, OH-VIN-1-3; *Microstegium vimineum* (Trin.) A. Camus, OH-VIN-2-1-S13, Vinton, Ohio, USA, 2020, WVA, Scott, OH-VIN-2-1; *Microstegium vimineum* (Trin.) A. Camus, PA-Berks-1940-233a-S37, Berks, Pennsylvania, USA, 1940, PH, Brumbach, 3277; *Microstegium vimineum* (Trin.) A. Camus, PA-Berks-1940-75-S36, Berks, Pennsylvania, USA, 1940, PH, Wilkens, 6471; *Microstegium vimineum* (Trin.) A. Camus, Bucks-1952-George-S45, Bucks, Pennsylvania, USA, 1952, PH, Long, 75812; *Microstegium vimineum* (Trin.) A. Camus, PA-Bucks-1954-26-S48, Bucks, Pennsylvania, USA, 1954, PH, Wherry, sn; *Microstegium vimineum* (Trin.) A. Camus, PA-Berks-1957b-S41, Berks, Pennsylvania, USA, 1957, PH, Wilkens, 9182; *Microstegium vimineum* (Trin.) A. Camus, PA-Berks-1959-S43, Berks, Pennsylvania, USA, 1959, PH, Berkheimer, 19765; *Microstegium vimineum* (Trin.) A. Camus, PA-Montgomery-1959-S42, Montgomery, Pennsylvania, USA, 1959, PH, Wherry, sn; *Microstegium vimineum* (Trin.) A. Camus, PA-Fulton-1995-79-S40, Fulton, Pennsylvania, USA, 1995, CM, Grund, 1392; *Microstegium vimineum* (Trin.) A. Camus, PA-ALLE-1-1-S7, Allegheny, Pennsylvania, USA, 2020, WVA, Kuebbing, PA-ALLE-1-1; *Microstegium vimineum* (Trin.) A. Camus, PA-BUT-1-1-S16, Butler, Pennsylvania, USA, 2020, WVA, Heberling, PA-BUT-1-1; *Microstegium vimineum* (Trin.) A. Camus, PA-MONT-1-1-S3, Montgomery, Pennsylvania, USA, 2020, WVA, Moore, PA-MONT-1-1; *Microstegium vimineum* (Trin.) A.

Camus, PR-ElVerdeExSta-1966-52-S23, Rio Grande, Puerto Rico, USA, 1966, BRIT, Duncan, sn; *Microstegium vimineum* (Trin.) A. Camus, TN-Anderson-1934-53-S24, Anderson, Tennessee, USA, 1934, TENN, Jennison, 4360; *Microstegium vimineum* (Trin.) A. Camus, TN-Anderson-1934-S61, Anderson, Tennessee, USA, 1934, BRIT, Jennison, 3348; *Microstegium vimineum* (Trin.) A. Camus, TN-Knox-1934-S65, Knox, Tennessee, USA, 1934, TENN, Miller, 3482; *Microstegium vimineum* (Trin.) A. Camus, TN-Knox-1936-16-S9, Knox, Tennessee, USA, 1936, MO, Jennison, 260; *Microstegium vimineum* (Trin.) A. Camus, TN-Roane-1956-S66, Roane, Tennessee, USA, 1956, TENN, Norris & DeSelm, 21779; *Microstegium vimineum* (Trin.) A. Camus, TN-Bount-1961-S62, Blount, Tennessee, USA, 1961, TENN, Pringle, 29862; *Microstegium vimineum* (Trin.) A. Camus, TN-Knox-1970-S74, Knox, Tennessee, USA, 1970, TENN, Somers & Bowers, 81; *Microstegium vimineum* (Trin.) A. Camus, TN-Roane-1974-80-S41, Roane, Tennessee, USA, 1974, CM, Hedge, 50096; *Microstegium vimineum* (Trin.) A. Camus, TN-Carrol-2007-S71, Carrol, Tennessee, USA, 2007, TENN, Crabtree & McCoy, sn; *Microstegium vimineum* (Trin.) A. Camus, TN-Cheatham-2010-S68, Cheatham, Tennessee, USA, 2010, TENN, Klagstad, 432; *Microstegium vimineum* (Trin.) A. Camus, TN-KNO-3C-2-1-S22, Knox, Tennessee, USA, 2020, WVA, Barrett, TN-KNO-3C-2-1-S22; *Microstegium vimineum* (Trin.) A. Camus, TN-KNO-3C-2-6-B-S44, Knox, Tennessee, USA, 2020, WVA, Barrett, TN-KNO-3C-2-6-B-S44; *Microstegium vimineum* (Trin.) A. Camus, TN-KNO-7I-2-1-B-S47, Knox, Tennessee, USA, 2020, WVA, Barrett, TN-KNO-7I-2-1-B-S47; *Microstegium vimineum* (Trin.) A. Camus, TN-KNO-HP-1-1-A-S24, Knox, Tennessee, USA, 2020, WVA, Barrett, TN-KNO-HP-1-1-A-S24; *Microstegium vimineum* (Trin.) A. Camus, TN-KNO-HP-1-1-B-S45, Knox, Tennessee, USA, 2020, WVA, Barrett, TN-KNO-HP-1-1-B-S45; *Microstegium vimineum* (Trin.) A. Camus, TN-KNO-HP-1-6-A-S25, Knox, Tennessee, USA, 2020, WVA, Barrett, TN-KNO-HP-1-6-A-S25; *Microstegium vimineum* (Trin.) A. Camus, TN-KNO-HP-1-6-B-S46, Knox, Tennessee, USA, 2020, WVA, Barrett, TN-KNO-HP-1-6-B-S46; *Microstegium vimineum* (Trin.) A. Camus, TN-LPG-1-1-S59, Greene, Tennessee, USA, 2020, WVA, Barrett, TN-LPG-1-1-S59; *Microstegium vimineum* (Trin.) A. Camus, VA-Fairfax-1986-S80, Fairfax, Virginia, USA, 1986, TENN, Fosberg, 65307; *Microstegium vimineum* (Trin.) A. Camus, 206-VA-Bath-1-4-S8, Bath, Virginia, USA, 2019, WVA, Barrett, VA-BATH-1-4; *Microstegium vimineum* (Trin.) A. Camus, VA-SG-2-1-S49, Smyth, Virginia, USA, 2020, WVA, Barrett, VA-SMYTH-2-1; *Microstegium vimineum* (Trin.) A. Camus, VA-SG-2-6-S50, Smyth, Virginia, USA, 2020, WVA, Barrett, VA-SMYTH-2-6; *Microstegium vimineum* (Trin.) A. Camus, WV-Fayette-1977-105-S54, Fayette, West Virginia, USA, 1977, WVA, Grafton, sn; *Microstegium vimineum* (Trin.) A. Camus, 104-Cabell-Boone-WV-1987-S7, Cabell, West Virginia, USA, 1987, WVA, Cusick, 27164; *Microstegium vimineum* (Trin.) A. Camus, WV-Fayette-1997-103-S53, Fayette, West Virginia, USA, 1997, WVA, Grafton, sn; *Microstegium vimineum* (Trin.) A. Camus, WV-Calhoun-2000-96-S52, Calhoun, West Virginia, USA, 2000, WVA, Grafton, sn; *Microstegium vimineum* (Trin.) A. Camus, 99-WV-Clay-2002-S6, Clay, West Virginia, USA, 2002, WVA, Grafton, sn; *Microstegium vimineum* (Trin.) A. Camus, WV-Hardy-2003-107-S56, Hardy, West Virginia, USA, 2003, WVA, Grafton, sn; *Microstegium vimineum* (Trin.) A. Camus, WV-Harrison-2007-106-S55, Harrison, West Virginia, USA, 2007, WVA, Grafton, sn; *Microstegium vimineum* (Trin.) A. Camus, WV-Marion-2007-145-S74, Marion, West Virginia, USA, 2007, WVA, Grafton, sn; *Microstegium vimineum* (Trin.) A. Camus, WV-SR-2-3-225-S85, Preston, West Virginia, USA, 2019, WVA, Huebner & Barrett, WV-SR-2-1; *Microstegium vimineum* (Trin.) A. Camus, WI-LAC-2-1-S8, LaCrosse, Wisconsin, USA, 2020, WVA, Molano-Flores, WI-LAC-2-1; *Microstegium vimineum* (Trin.) A. Camus, WI-LAC-2-2-S9, LaCrosse, Wisconsin, USA, 2020, WVA, Molano-Flores, WI-LAC-2-2.

## Literature Cited

Adobe Inc. 2019 Adobe Illustrator. Retrieved from https://adobe.com/products/illustrator.

Alverson AJ, DW Rice, S Dickinson, K Barry, JD Palmer 2011 Origins and recombination of the bacterial-sized multichromosomal mitochondrial genome of cucumber. Plant Cell 23:2499–2513.

Alverson AJ, X Wei, DW Rice, DB Stern, K Barry, JD Palmer 2010 Insights into the evolution of mitochondrial genome size from complete sequences of *Citrullus lanatus* and *Cucurbita pepo* (Cucurbitaceae). Mol Biol Evol 27:1436–1448.

Amos B, C Aurrecoechea, M Barba, A Barreto, EY Basenko, W Bażant, R Belnap, AS Blevins, U Böhme, J Brestelli, et al. 2022 VEuPathDB: the eukaryotic pathogen, vector and host bioinformatics resource center. Nucleic Acids Res 50:D898–D911.

Arthan W, MR McKain, P Traiperm, CAD Welker, JK Teisher, EA Kellogg 2017 Phylogenomics of Andropogoneae (Panicoideae: Poaceae) of Mainland Southeast Asia. Syst Bot 42:418–431.

Banerjee AK, W Guo, Y Huang 2019 Genetic and epigenetic regulation of phenotypic variation in invasive plants – linking research trends towards a unified framework. NeoBiota 49:77–103.

Bao W, KK Kojima, O Kohany 2015 Repbase Update, a database of repetitive elements in eukaryotic genomes. Mobile DNA 6:11.

Barrett CF, WJ Baker, JR Comer, JG Conran, SC Lahmeyer, JH Leebens Mack, J Li, GS Lim, DR Mayfield Jones, L Perez, et al. 2016 Plastid genomes reveal support for deep phylogenetic relationships and extensive rate variation among palms and other commelinid monocots. New Phytol 209:855–870.

Barrett CF, CD Huebner, ZA Bender, TA Budinsky, CW Corbett, M Latvis, MR McKain, M Motley, SV Skibicki, HL Thixton, et al. 2022 Digitized collections elucidate invasion history and patterns of awn polymorphism in *Microstegium vimineum*. Am J Bot 109:689–705.

Bendich AJ 1993 Reaching for the ring: the study of mitochondrial genome structure. Curr Genet 24:279–290.

Bieker VC, P Battlay, B Petersen, X Sun, J Wilson, JC Brealey, F Bretagnolle, K Nurkowski, C Lee, FS Barreiro, et al. 2022 Uncovering the genomic basis of an extraordinary plant invasion. Sci Adv 8:eabo5115.

van Boheemen LA, E Lombaert, KA Nurkowski, B Gauffre, LH Rieseberg, KA Hodgins 2017 Multiple introductions, admixture and bridgehead invasion characterize the introduction history of *Ambrosia artemisiifolia* in Europe and Australia. Mol Ecol 26:5421–5434.

Camacho C, G Coulouris, V Avagyan, N Ma, J Papadopoulos, K Bealer, TL Madden 2009 BLAST+: architecture and applications. BMC Bioinform 10:421.

Cardi T, P Giegé, S Kahlau, N Scotti 2012 Expression profiling of organellar genes. In: Bock R, Knoop V, editors. Genomics of chloroplasts and mitochondria. Pages 323–355 *In* Advances in photosynthesis and respiration. Vol. 35. Springer Netherlands, Dordrecht.

Cavanagh AM, JW Morgan, RC Godfree 2020 Awn morphology influences dispersal, microsite selection and burial of Australian native grass diaspores. Front Ecol Evol 8. [accessed 2023 Feb 7]. https://www.frontiersin.org/articles/10.3389/fevo.2020.581967

Chen C-H, JF Veldkamp, C-S Kuoh, C-C Tsai, Y-C Chiang 2009 Segregation of *Leptatherum* from *Microstegium* (Andropogoneae, Poaceae) confirmed by Internal Transcribed Spacer DNA sequences. Blumea 54:175–180.

Chen C-H, J-F Veldkamp, C-S Kuoh 2012 Taxonomic revision of *Microstegium* s.str. (Andropogoneae, Poaceae). Blumea 57:160–189.

Clifton SW, P Minx, CM-R Fauron, M Gibson, JO Allen, H Sun, M Thompson, WB Barbazuk, S Kanuganti, C Tayloe, et al. 2004 Sequence and comparative analysis of the maize NB mitochondrial genome. Plant Physiol 136:3486–3503.

Crotty SM, BQ Minh, NG Bean, BR Holland, J Tuke, LS Jermiin, AV Haeseler 2020 GHOST: Recovering historical signal from heterotachously evolved sequence alignments. Syst Biol 69:249–264.

Daehler CC 1998 The taxonomic distribution of invasive angiosperm plants: Ecological insights and comparison to agricultural weeds. Biol Conserv 84:167–180.

Dlugosch KM, IM Parker 2008 Founding events in species invasions: genetic variation, adaptive evolution, and the role of multiple introductions. Mol Ecol 17:431–449.

Dlugosch KM, FA Cang, BS Barker, K Andonian, SM Swope, LH Rieseberg 2015 Evolution of invasiveness through increased resource use in a vacant niche. Nat Plants 1:15066.

Doyle JJ, JL Doyle 1987 A rapid DNA isolation procedure for small quantities of fresh leaf tissue. Phytochemical Bulletin, 19:11–15.

Estep MC, MR McKain, D Vela Diaz, J Zhong, JG Hodge, TR Hodkinson, DJ Layton, ST Malcomber, R Pasquet, EA Kellogg 2014 Allopolyploidy, diversification, and the Miocene grassland expansion. PNAS 111:15149–15154.

Fairbrothers DE, JR Gray 1972 *Microstegium vimineum* (Trin.) A. Camus (Gramineae) in the United States. J Torrey Bot 99:97–100.

Fang Y, H Wu, T Zhang, M Yang, Y Yin, L Pan, et al. 2012 A complete sequence and transcriptomic analyses of date palm (*Phoenix dactylifera L*.) mitochondrial genome. PLoS ONE 7: e37164.

Finch DM, JL Butler, JB Runyon, CJ Fettig, FF Kilkenny, S Jose, SJ Frankel, SA Cushman, RC Cobb, JS Dukes, et al. 2021 Effects of climate change on invasive species. Pages 57–83. In Poland TM, Patel-Weynand T, Finch DM, Miniat CF, Hayes DC, Lopez VM, eds. Invasive species in forests and rangelands of the United States. Cham: Springer International Publishing. [accessed 2023 Feb 1]. https://link.springer.com/10.1007/978-3-030-45367-1_4

Flory SL, F Long, K Clay 2011 Invasive *Microstegium* populations consistently outperform native range populations across diverse environments. Ecology 92:2248–2257.

Frankham R 2005 Resolving the genetic paradox in invasive species. Heredity 94:385–385.

Givnish TJ, M Ames, JR McNeal, MR McKain, PR Steele, CW dePamphilis, SW Graham, JC Pires, DW Stevenson, WB Zomlefer, et al. 2010 Assembling the Tree of the Monocotyledons: plastome sequence phylogeny and evolution of Poales. Ann Missouri Bot Gard 97:584–616.

Gualberto JM, D Mileshina, C Wallet, AK Niazi, F Weber-Lotfi, A Dietrich 2014 The plant mitochondrial genome: dynamics and maintenance. Biochimie 100:107–120.

Harris RS 2007 Improved pairwise alignment of genomic DNA. Ph.D. Thesis, The Pennsylvania State University.

Hawkins JS, D Ramachandran, A Henderson, J Freeman, M Carlise, A Harris, Z Willison-Headley 2015 Phylogenetic reconstruction using four low-copy nuclear loci strongly supports a polyphyletic origin of the genus *Sorghum*. Ann Bot 116:291–299.

Hoang DT, O Chernomor, A von Haeseler, BQ Minh, LS Vinh 2018 UFBoot2: Improving the ultrafast bootstrap approximation. Mol Biol and Evol 35:518–522.

Hodgins KA, L Rieseberg, SP Otto 2009 Genetic control of invasive plants species using selfish genetic elements. Evol Appl 2:555–569.

Holec S, H Lange, K Kühn, M Alioua, T Börner, D Gagliardi 2006 Relaxed transcription in *Arabidopsis* mitochondria is counterbalanced by RNA stability control mediated by polyadenylation and polynucleotide phosphorylase. Mol Cell Biol 26:2869–2876.

Hu J, Y Zheng, X Shang 2018 MiteFinderII: a novel tool to identify miniature inverted-repeat transposable elements hidden in eukaryotic genomes. BMC Medical Genom 11:101.

Hudson J, JC Castilla, PR Teske, LB Beheregaray, ID Haigh, CD McQuaid, M Rius 2021 Genomics-informed models reveal extensive stretches of coastline under threat by an ecologically dominant invasive species. PNAS 118:e2022169118.

Huebner CD 2010a Establishment of an invasive grass in closed-canopy deciduous forests across local and regional environmental gradients. Biol Invasions 12:2069–2080.

Huebner CD 2010b Spread of an invasive grass in closed-canopy deciduous forests across local and regional environmental gradients. Biol Invasions 12:2081–2089.

Huebner CD, CW Cameron Corbett, L Ferrari, LE Kosslow, SV Skibicki, CF Barrett 2022 Traits that define invasiveness in *Microstegium vimineum* (Japanese stiltgrass) at local and regional scales. Botany 2022, Anchorage, Alaska, United States.

Hunt M, ND Silva, TD Otto, J Parkhill, JA Keane, SR Harris 2015 Circlator: automated circularization of genome assemblies using long sequencing reads. Genome Biol 16:294.

Jackman SD, L Coombe, RL Warren, H Kirk, E Trinh, T MacLeod, S Pleasance, P Pandoh, Y Zhao, RJ Coope, et al. 2020 Complete mitochondrial genome of a gymnosperm, sitka spruce (*Picea sitchensis*), indicates a complex physical structure. Genome Biol Evol 12:1174–1179.

Johnson DJ, SL Flory, A Shelton, C Huebner, K Clay 2015 Interactive effects of a non-native invasive grass *Microstegium vimineum* and herbivore exclusion on experimental tree regeneration under differing forest management. J Appl Ecol 52:210–219.

Kalyaanamoorthy S, BQ Minh, TKF Wong, A von Haeseler, LS Jermiin 2017 ModelFinder: fast model selection for accurate phylogenetic estimates. Nat Methods 14:587–589.

Kassambara A 2020 ggpubr: ‘ggplot2’ based publication ready plots. R package version 0.4.0. https://CRAN.R-project.org/package=ggpubr.

Keller SR, DR Taylor 2010 Genomic admixture increases fitness during a biological invasion. J Evol Biol 23:1720–1731.

Kerns BK, C Tortorelli, MA Day, T Nietupski, AMG Barros, JB Kim, MA Krawchuk 2020 Invasive grasses: A new perfect storm for forested ecosystems? For Ecol Manag 463:117985.

Knoop V, M Unseld, J Marienfeld, P Brandt, S Sünkel, H Ullrich, A Brennicke 1996 Copia-, Gypsy- and LINE-Like retrotransposon fragments in the mitochondrial genome of *Arabidopsis thaliana*. Genetics 142:579–585.

Kolbe JJ, RE Glor, L Rodríguez Schettino, AC Lara, A Larson, JB Losos 2004 Genetic variation increases during biological invasion by a Cuban lizard. Nature 431:177–181.

Kolmogorov M, J Yuan, Y Lin, PA Pevzner 2019 Assembly of long, error-prone reads using repeat graphs. Nat Biotechnol 37:540–546.

Koren S, BP Walenz, K Berlin, JR Miller, NH Bergman, AM Phillippy 2017 Canu: scalable and accurate long-read assembly via adaptive *k*-mer weighting and repeat separation. Genome Res 27:722–736.

Kovar L, M Nageswara-Rao, S Ortega-Rodriguez, DV Dugas, S Straub, R Cronn, SR Strickler, CE Hughes, KA Hanley, DN Rodriguez, et al. 2018 PacBio-based mitochondrial genome assembly of *Leucaena trichandra* (Leguminosae) and an intrageneric assessment of mitochondrial RNA editing. Genome Biol Evol 10:2501–2517.

Kurtz S 2001 REPuter: the manifold applications of repeat analysis on a genomic scale. Nucleic Acids Res 29:4633–4642.

Lehwark P, S Greiner 2019 GB2sequin - A file converter preparing custom GenBank files for database submission. Genomics 111:759–761.

Li H, B Handsaker, A Wysoker, T Fennell, J Ruan, N Homer, G Marth, G Abecasis, R Durbin, 1000 Genome Project Data Processing Subgroup 2009 The sequence alignment/map format and SAMtools. Bioinformatics 25:2078–2079.

Lin Y, P Li, Y Zhang, D Akhter, R Pan, Z Fu, M Huang, X Li, Y Feng 2022 Unprecedented organelle genomic variations in morning glories reveal independent evolutionary scenarios of parasitic plants and the diversification of plant mitochondrial complexes. BMC Biol 20:49.

Lloyd Evans D, SV Joshi, J Wang 2019 Whole chloroplast genome and gene locus phylogenies reveal the taxonomic placement and relationship of *Tripidium* (Panicoideae: Andropogoneae) to sugarcane. BMC Evol Biol 19:33.

Ma J, JL Bennetzen 2004 Rapid recent growth and divergence of rice nuclear genomes. PNAS 101:12404–12410.

Maier RM, P Zeitz, H Kössel, G Bonnard, JM Gualberto, JM Grienenberger 1996 RNA editing in plant mitochondria and chloroplasts. Plant Mol Biol 1996 32:343–65

Marienfeld J, M Unseld, A Brennicke 1999 The mitochondrial genome of *Arabidopsis* is composed of both native and immigrant information. Trends in Plant Sci 4:495–502.

Matheson P, A McGaughran 2022 Genomic data is missing for many highly invasive species, restricting our preparedness for escalating incursion rates. Sci Rep 12:13987.

Mehrhoff LJ 2000 Perennial *Microstegium vimineum* (Poaceae): An apparent misidentification? J Torrey Bot Soc 127:251.

Minh BQ, HA Schmidt, O Chernomor, D Schrempf, MD Woodhams, A von Haeseler, R Lanfear 2020 IQ-TREE 2: New models and efficient methods for phylogenetic inference in the genomic era. Teeling E, editor. Mol Biol Evol 37:1530–1534.

Mortensen DA, ESJ Rauschert, AN Nord, BP Jones 2009 Forest roads facilitate the spread of invasive plants. Invasive Plant Cci Manag 2:191–199.

Mounger J, ML Ainouche, O Bossdorf, A Cavé-Radet, B Li, M Parepa, A Salmon, J Yang, CL Richards Epigenetics and the success of invasive plants. Philos Trans R Soc Lond B Biol Sci 376:20200117.

Mower JP, DB Sloan, AJ Alverson 2012 Plant mitochondrial genome diversity: the genomics revolution. Pages 123–144 In Wendel JF, Greilhuber J, Dolezel J, Leitch IJ, editors. Plant Genome Diversity Volume 1: Plant genomes, their residents, and their evolutionary dynamics. Springer, Vienna.

North HL, A McGaughran, CD Jiggins 2021 Insights into invasive species from whole-genome resequencing. Mol Ecol 30:6289–6308.

Notsu Y, S Masood, T Nishikawa, N Kubo, G Akiduki, M Nakazono, A Hirai, K Kadowaki 2002 The complete sequence of the rice (*Oryza sativa* L.) mitochondrial genome: frequent DNA sequence acquisition and loss during the evolution of flowering plants. Mol Gen Genomics 268:434–445.

Novy A, S l. Flory, JM Hartman 2013 Evidence for rapid evolution of phenology in an invasive grass. J Evol Biol 26:443–450.

Ortiz EM 2019 vcf2phylip v2.0: convert a VCF matrix into several matrix formats for phylogenetic analysis. DOI:10.5281/zenodo.2540861

Palmer JD, LA Herbon 1988 Plant mitochondrial DNA evolves rapidly in structure, but slowly in sequence. J Mol Evol 28:87–97.

Pimentel D, R Zuniga, D Morrison 2005 Update on the environmental and economic costs associated with alien-invasive species in the United States. Ecol Econ 52:273–288.

Poplin R, V Ruano-Rubio, MA DePristo, TJ Fennell, MO Carneiro, GA Van der Auwera, DE Kling, LD Gauthier, A Levy-Moonshine, D Roazen, et al. 2017 Scaling accurate genetic variant discovery to tens of thousands of samples. bioRxiv:201178.

Ramachandran D, CD Huebner, M Daly, J Haimovitz, T Swale, CF Barrett 2021 chromosome level genome assembly and annotation of highly invasive Japanese stiltgrass (*Microstegium vimineum*). Genome Biol Evol 13:evab238.

Ranwez V, EJP Douzery, C Cambon, N Chantret, F Delsuc 2018 MACSE v2: Toolkit for the alignment of coding sequences accounting for frameshifts and stop codons. Mol Biol Evol 35:2582–2584.

Rauschert ESJ, DA Mortensen, ON Bjørnstad, AN Nord, N Peskin 2010 Slow spread of the aggressive invader, *Microstegium vimineum* (Japanese stiltgrass). Biol Invasions 12:563– 579.

Revolinski SR, PJ Maughan, CE Coleman, IC Burke 2022 Preadapted to adapt: underpinnings of adaptive plasticity revealed by the downy brome genome. In Review. [accessed 2023 Feb 1]. https://www.researchsquare.com/article/rs-2050485/v1

Rice DW, AJ Alverson, AO Richardson, GJ Young, MV Sanchez-Puerta, J Munzinger, K Barry, JL Boore, Y Zhang, CW dePamphilis, et al. 2013 Horizontal transfer of entire genomes via mitochondrial fusion in the angiosperm *Amborella*. Science 342:1468–1473.

Rius M, JA Darling 2014 How important is intraspecific genetic admixture to the success of colonising populations? Trends Ecol Evol 29:233–242.

Ruwe H, G Wang, S Gusewski, C Schmitz-Linneweber 2016 Systematic analysis of plant mitochondrial and chloroplast small RNAs suggests organelle-specific mRNA stabilization mechanisms. Nucleic Acids Res 44:7406–7417.

Sakai AK, FW Allendorf, JS Holt, DM Lodge, J Molofsky, KA With, S Baughman, RJ Cabin, JE Cohen, NC Ellstrand, et al. 2001 The population biology of invasive species. Ann Rev of Ecol Systematics 32:305–332.

Sanchez Puerta MV, LE García, J Wohlfeiler, LF Ceriotti 2017 Unparalleled replacement of native mitochondrial genes by foreign homologs in a holoparasitic plant. New Phytol 214:376–387.

Simberloff D, J-L Martin, P Genovesi, V Maris, DA Wardle, J Aronson, F Courchamp, B Galil, E García-Berthou, M Pascal, et al. 2013 Impacts of biological invasions: what’s what and the way forward. Trends Ecol Evol 28:58–66.

Sinn BT, CF Barrett 2020 Ancient mitochondrial gene transfer between fungi and the orchids. Mol Biol Evol 37:44–57.

Sinn BT, SJ Simon, MV Santee, SP DiFazio, NM Fama, CF Barrett 2022 ISSRseq: An extensible method for reduced representation sequencing. Methods Ecol Evol, 13:668– 681.

Sloan DB 2013 One ring to rule them all? Genome sequencing provides new insights into the ‘master circle’ model of plant mitochondrial DNA structure. New Phytol 200:978–985.

Sloan DB, Z Wu 2014 History of plastid DNA insertions reveals weak deletion and AT mutation biases in Angiosperm mitochondrial genomes. Genome Biol Evol 6:3210–3221.

Smit AFA, R Hubley, P Green 2013 RepeatMasker Open-4.0. http://www.repeatmasker.org.

Sutherland BL, CF Barrett, JB Beck, M Latvis, MR McKain, EM Sigel, NJ Kooyers 2021 Botany is the root and the future of invasion biology. Am J Bot 108:549–552.

Turner KG, KL Ostevik, CJ Grassa, LH Rieseberg 2021 Genomic analyses of phenotypic differences between native and invasive populations of diffuse knapweed (Centaurea diffusa). Frontiers Ecol Evol 8. [accessed 2023 Feb 1]. https://www.frontiersin.org/articles/10.3389/fevo.2020.577635

Van der Auwera GA, Carneiro M, Hartl C, Poplin R, del Angel G, Levy-Moonshine A, Jordan T, Shakir K, Roazen D, Thibault J, et al. 2013 From FastQ data to high-confidence variant calls: the genome analysis toolkit best practices pipeline. Curr Protoc Bioinformatics, 43:11.10.1-11.10.33.

Van der Auwera GA, BD O’Connor 2020 Genomics in the Cloud: Using Docker, GATK, and WDL in Terra (1st Edition). O’Reilly Media.

Verhoeven KJF, M Macel, LM Wolfe, A Biere 2010 Population admixture, biological invasions and the balance between local adaptation and inbreeding depression. Proc Royal Soc B 278:2–8.

Walker BJ, T Abeel, T Shea, M Priest, A Abouelliel, S Sakthikumar, CA Cuomo, Q Zeng, J Wortman, SK Young, et al. 2014 Pilon: An integrated tool for comprehensive microbial variant detection and genome assembly improvement. Wang J, editor. PLoS ONE 9:e112963.

Watson L, Dallwitz MJ 1994 The grass genera of the world. CAB International, Wallingford, Oxfordshire, UK.

Weiher E, PA Keddy 1995 The assembly of experimental wetland plant communities. Oikos 73:323–335.

Welker CAD, MR McKain, MC Estep, RS Pasquet, G Chipabika, B Pallangyo, EA Kellogg 2020 Phylogenomics enables biogeographic analysis and a new subtribal classification of Andropogoneae (Poaceae—Panicoideae). J Systematics Evol 58:1003–1030.

Wick RR, MB Schultz, J Zobel, KE Holt 2015 Bandage: interactive visualization of de novo genome assemblies. Bioinformatics 31:3350–3352.

Wickham H 2016 ggplot2: elegant graphics for data analysis. Springer-Verlag New York. ISBN 978–3-319-24277-4, https://ggplot2.tidyverse.org.

Wickham H, R François, L Henry, K Müller, D Vaughan 2023 dplyr: A grammar of data manipulation. https://dplyr.tidyverse.org, https://github.com/tidyverse/dplyr

Wood DE, J Lu, B Langmead 2019 Improved metagenomic analysis with Kraken 2. Genome Biol 20:257.

Wu Z, JM Cuthbert, DR Taylor, DB Sloan 2015 The massive mitochondrial genome of the angiosperm *Silene noctiflora* is evolving by gain or loss of entire chromosomes. PNAS 112:10185–10191.

Wu Z, X Liao, X Zhang, LR Tembrock, A Broz 2022 Genomic architectural variation of plant mitochondria—A review of multichromosomal structuring. J Sytematics Evol 60:160– 168.

Xiong W, L He, J Lai, HK Dooner, C Du 2014 HelitronScanner uncovers a large overlooked cache of Helitron transposons in many plant genomes. PNAS 111:10263–10268.

Xiong Yanli, Q Yu, Yi Xiong, J Zhao, X Lei, L Liu, W Liu, Y Peng, J Zhang, D Li, et al. 2022 The complete mitogenome of Elymus sibiricus and insights into its evolutionary pattern based on simple repeat sequences of seed plant mitogenomes. Frontiers Plant Sci 12. [accessed 2023 Feb 1]. https://www.frontiersin.org/articles/10.3389/fpls.2021.802321

Yu X, W Jiang, W Tan, X Zhang, X Tian 2020 Deciphering the organelle genomes and transcriptomes of a common ornamental plant *Ligustrum quihoui* reveals multiple fragments of transposable elements in the mitogenome. Int J Biol Macromolecules 165:1988–1999.

Zhao N, Y Wang, J Hua 2018 the roles of mitochondrion in intergenomic gene transfer in plants: a source and a pool. Int J Mol Sci 19:547.

