## Supplementary figures and images for "Mitochondrial genome sequencing and analysis of the invasive *Microstegium vimineum*: a resource for systematics, invasion history, and management"

### Fig. S1

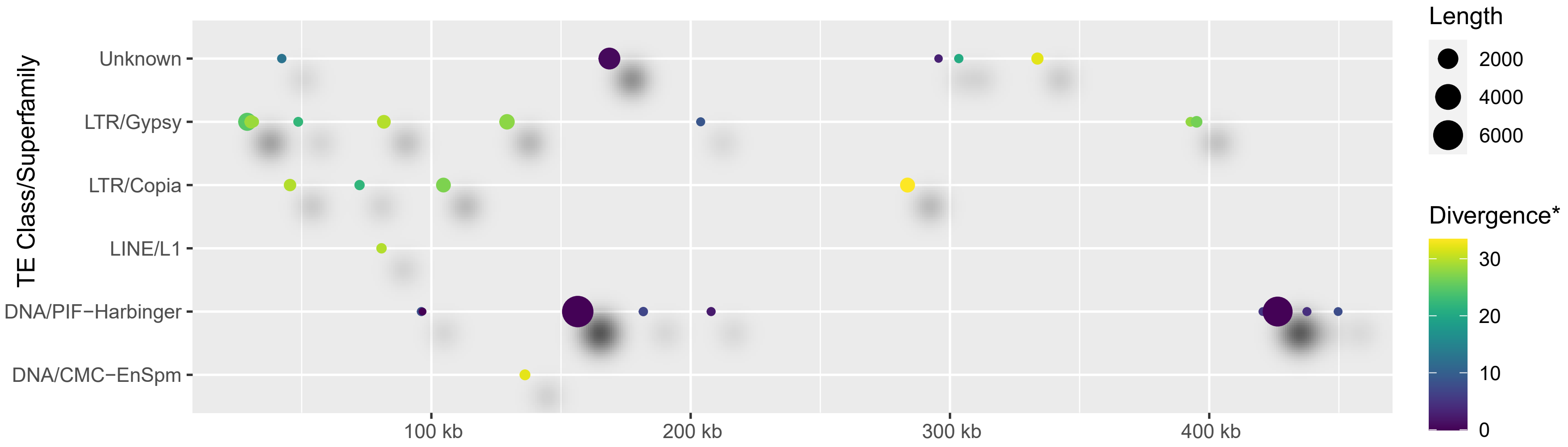

### Fig. S2

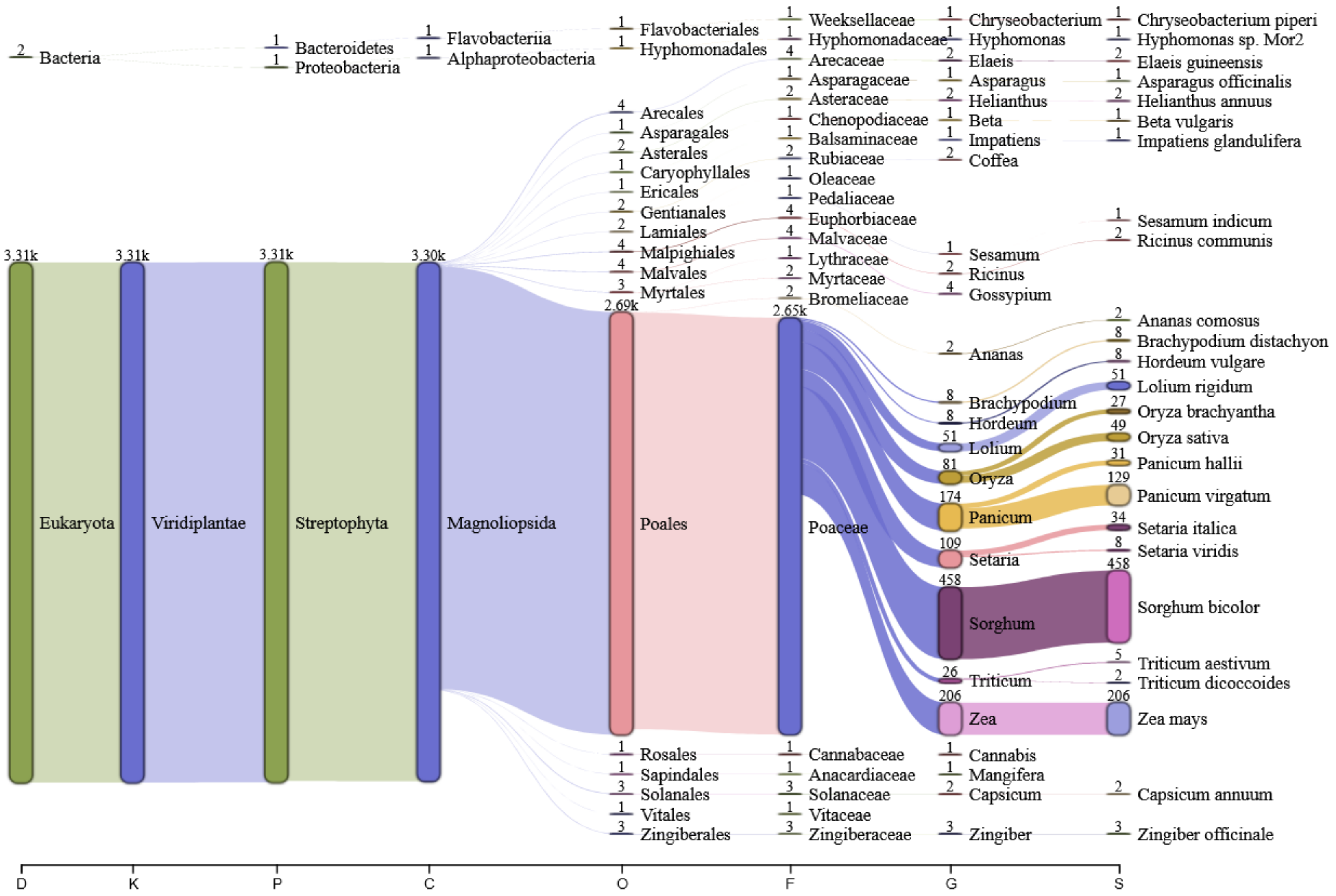
